# A genome-wide study of ruminants reveals two endogenous retrovirus families still active in goats

**DOI:** 10.1101/2024.06.21.600049

**Authors:** Marie Verneret, Caroline Leroux, Thomas Faraut, Vincent Navratil, Emmanuelle Lerat, Jocelyn Turpin

## Abstract

**Background:** Endogenous retroviruses (ERV) are traces of ancestral retroviral germline infections that constitute a significant portion of mammalian genomes and function as LTR-retrotransposons. ERVs remain understudied in ruminants and usually focus on specific families, highlighting a need for comprehensive and thorough exploration of the ERV landscape in these species.

**Results:** We characterized 23 *de novo* Class I and II ERV families across four reference assemblies of domestic and wild sheep and goats, and one assembly of cattle. Among these families, 15 were shared by the five ruminant species, while six were exclusive to small ruminants and two were exclusive to cattle. The presence of these families in other ruminant species revealed multiple endogenization events over 40 million years. We identified a total of 113,812 ERV insertions across the five genomes, representing between 0.5 and 1% of their genomes. Solo-LTRs account for 84.5% of the annotated copies demonstrating that most of the ERVs are relics of past events. Although Class I ERVs showed similar sequence divergence landscapes between species, Class II ERVs displayed contrasting evolutionary dynamics. Families II-3 and II-5 showed higher abundance and copy conservation in small ruminants. Family II-5 was closely related to circulating exogenous retroviruses and was identified with 22 copies sharing identical LTRs and 12 with complete coding capacities in the domestic goat.

**Conclusions:** Our results indicate that two ERV families may have retain their transpositional activity within ruminants, particularly in domestic goats, illustrating distinct evolutionary dynamics among ruminant species. This work highlights the ongoing influence of ERVs on genomic landscapes and provides further insights into their evolutionary trajectories in ruminant genomes.

## Background

Endogenous retroviruses (ERVs) are remnants of ancient retroviral germline infections that have become permanently integrated into the host genome and are transmitted vertically to subsequent generations the same way as the other host genes (1–3). These ERVs make up a large proportion of mammalian genomes, representing 8% and 10% of the human (4,5) and mouse genomes respectively (6). The genomic structure of ERVs consists of four retroviral genes —*gag*, *pro*, *pol*, and *env*— flanked by two long terminal repeats (LTRs) (7). They are considered as LTR-retrotransposons and can be classified into three groups according to the transposable element’s classification (8,9): ERV-1, ERV-K or ERV-L, or Class I, II or III according to the retroviral taxonomy (7) based on their relationship with the exogenous gammaretrovirus, betaretrovirus and spumaretrovirus genera respectively.

After their integration, most of the ERV insertions are subject to the effects of the genetic drift and become non-functional insertions due to the accumulation of point mutations, insertions, deletions, or even the complete excision of internal retroviral genes by recombination between the two LTRs, ultimately resulting in the formation of solo-LTR insertions (10–12). However, some ERV insertions have conserved their coding capacities and are then potentially capable to transpose through the copy-and-paste mechanism or capable of reinfecting other cells (13,14). ERVs may also be functionally relevant to their host organism (2). Some ERVs play a critical role in embryonic development, as exemplified by the syncytin gene, which has been captured and domesticated from a retroviral envelope several times in different mammalian species and is essential for placental development (15). In addition, LTR sequences can serve as a source of regulatory activity and affect the expression of cellular genes (16). However, ERVs can also have detrimental effects. Dysregulated transcription of ERVs has been linked to genome instability and has been implicated in a range of human diseases, including several cancers (17–19), neurodegenerative disorders (20,21), and autoimmune diseases (22–24).

While ERVs have been extensively studied in humans (25), the exploration of the ERV landscape in livestock organisms remains relatively understudied. ERVs are usually not directly related to existing circulating exogenous retroviruses. Small ruminants (including sheep and goats), along with koalas (26,27) and a few other vertebrate hosts (28,29), provide a unique paradigm with the coexistence of both endogenous retroviruses and their exogenous counterparts. In small ruminants, the exogenous Jaagsiekte sheep retrovirus (JSRV) and enzootic nasal tumor virus (ENTV), responsible for respiratory cancers (30–32), are closely related to an ERV family previously named enJSRV (33). Most studies on ERVs in small ruminants have focused on this family. Collectively, it has been shown that these ERVs have evolved over millions of years in the sheep (34,35). They play a critical role in sheep reproductive physiology (36,37) and also appear to confer potential host protection against exogenous retrovirus infection (38–40) although the extent of this protection *in vivo* still remains unclear (41). Other studies have shown that enJSRV insertions can have phenotypic effects (42) and are also highly polymorphic in sheep populations (43).

However, despite the focus on the enJSRV family, the global ERV landscape of small ruminants remains uncharacterized, and the function of the other families is largely unknown. This study explores this unique model using high-resolution characterization techniques and bioinformatic analyses to unravel the comprehensive ERV repertoire in ruminant genomes and assess their past and present evolutionary history. Through this initiative, we aim to elucidate the complex interplay between ERVs and their host genomes, providing insights into their role in shaping genome diversity and adaptation in small ruminants.

## Methods

### *De novo* characterization of ERV consensus sequences

Endogenous retroviruses (ERVs) have been characterized in five ruminant species: cattle (*Bos taurus*), domestic sheep (*Ovis aries*), wild sheep (*Ovis orientalis*), domestic goat (*Capra hircus*), and wild goat (*Capra aegagrus*). These five species were specifically selected because of the high quality of their assemblies available at the chromosome level and serving as reference genome in their respective species. Each assembly was retrieved from the Genbank database (accession numbers in Table S1) and ERV consensus sequences were identified using RepeatModeler (version 2.0.3) (44) with default parameters (Fig. S1).

The raw consensus sequences identified by RepeatModeler, were filtered out with an approach inspired from Goubert *et al* (45). Briefly, only sequences classified as ERV were conserved. The ones ranging from 1 to 15 kb were considered as potential internal parts (INT) with the retroviral genes and those smaller than 1 kb as long terminal repeats (LTRs). According to the 80-80-80 rule (8), sequences longer than 80bp that share more than 80% identity over 80% of their sequences belong to the same family. To reduce redundancy, INT and LTR sequences were clustered separately using CD-HIT (46). When sequences had more than 80% of sequence identity, only one representative sequence was kept for each cluster.

The retroviral genes (*gag*, *pro*/*pol*, and *env*) were annotated within each INT consensus sequence using BLASTx (47) against a subset of the Dfam database (version 3.7) (48), that we named ‘dfam_retro’, obtained using the keywords “gag”, “pro”, “pol”, “env”. To minimize false positives, only INT consensus sequences with at least one hit on a retroviral gene from “dfam_retro” were conserved. The different reading frames were also identified using ORFFinder (https://www.ncbi.nlm.nih.gov/orffinder/) (49). To re-associate the INT and LTR consensus sequences, the filtered INT and LTR consensus sequences were used as a custom library in RepeatMasker (version 4.1.5) (50) to identify hits corresponding to ERV copies in the five assemblies. Copies were reconstructed when INT and LTR hits were located within 500bp from each other. Consensus sequences representing less than 10 putative copies were removed and the association between INT and LTR consensus sequences was established when they were connected in at least 10 copies. When the internal consensus parts were not associated with any LTR consensus sequence, the true absence of LTR was verified by examining the corresponding annotated copies. The position of these copies was extended by 5kb on both the 5’ and 3’ sides and extracted from the genome using BEDTools (version 2.30.0) (51). A multiple sequence alignment of the extended copies was performed using MAFFT (version 7.490) (52) and visualized using Geneious Prime 2023.0.2 (https://www.geneious.com). The start and the end of the copies were determined by identifying blocks of homology and the flanking sequences were manually cropped. The presence of flanking LTRs was determined using self dotplot. When LTR sequences were identified, LTR consensus sequences were reconstructed using all the LTR sequences from the copies. The final set of consensus sequences is available as supplementary data (Files S1 to S5).

### Phylogenetic analyses and classification of the consensus sequences

The characterized consensus sequences were aligned with retroviral sequences from Genbank including gammaretroviruses such as Gibbon Ape Leukemia Virus (GaLV: NC_001885.3), Koala Retrovirus (KoRV: NC_039228.1), Murine Leukemia Virus (MuLV: NC_001501.1), Feline Leukemia Virus (FeLV: AF052723.1), as well as betaretroviruses with Mouse Mammary Tumor Virus (MMTV: NC_001503.1), Enzootic Nasal Tumor Virus Type 1 and 2 (ENTV1: NC_007015.1 and ENTV2: NC_004994.2), Jaagsiekte Sheep Retrovirus (JSRV: AF105220.1). Endogenous retrovirus sequences including human endogenous retroviruses from group E, K and L (HERV-E : M10976.1, HERV-K : M14123.1, HERV-L : X89211.1), feline endogenous retrovirus (enFeLV: AY364318.1), murine endogenous retrovirus (MuERV-L: Y12713.1), and cattle and sheep ERV reference consensus from Repbase (version 29.01) (53) were also included. The references available in Repbase for *Capra aegagrus* were not included as they contain only LTR sequences and no retroviral genes. Alignment was performed using MAFFT (52) followed by phylogenetic analysis using Maximum Likelihood (ML) statistical method in IQ-TREE (version 1.5.3) (54,55) with 10,000 ultrafast bootstrap replicates (56). Phylogenetic trees were visualized using the online version of the Interactive Tree Of Life (iTOL, https://itol.embl.de) (57). Finally, each ERV family was categorized as Class I or Class II ERVs based on the closest exogenous retroviruses respectively gamma or betaretroviruses. The name of the ERV families were chosen according to their classification as Class I or II along with an arbitrary attributed number.

### ERV annotation in reference assemblies

The characterized consensus sequences were used as a custom library to annotate the ovine, caprine and bovine reference assemblies using RepeatMasker (version 4.1.5) (50). Hits that shared at least 80% of sequence identity with the corresponding consensus sequence and were greater than 80 bp were conserved (8). Features originating from the same family and located closer than 500bp for solo-LTRs and 7kb for other copies were merged to reconstruct the different ERV insertions. The complete annotation of the five selected assemblies and the number of insertions per family are available as supplementary data (Files S6-S10 and Table S3). The ERV genome fraction was computed as the proportion of bases in the genome covered by all the detected ERV insertions (ERV number of bases / total genome length x 100). Each copy was aligned to its consensus sequence using MAFFT (52) and a divergence score was calculated using the Kimura-2-parameter model (K80) with the ape package in R (version 4.2.3) (58,59). In cases where an insertion had multiple hits on different consensus sequences, the sub-family with the lowest divergence score was assigned. The metadata associated with the insertions are provided as supplementary data (Files S11-S15).

The 5’ and 3’ LTR sequences were extracted from each ERV insertions and pairwise aligned using MAFFT (52) to estimate their sequence divergence. The retroviral open reading frames (ORF) in the ERV insertions were annotated using orfipy (60). Only ORFs starting with an ATG codon and having a minimum length of 300bp were conserved for *gag* and *env*. For *pro* and *pol*, as they are produced by frameshifts, ORFs with any sense codon longer than 300bp were retained. Each ORF was translated into a protein sequence and aligned against the “dfam_retro” database (see above) using BLASTp (47). Insertions with intact *gag*, *pro/pol* and *env* ORFs (>80% of the expected length) were designated as full-length copies potentially capable of reinfecting other cells. Those lacking only the *env* ORF were considered as copies capable of retrotransposition (Table S3).

### ERV insertion sites comparison

A comparative analysis of the families II-3 and II-5 insertion sites in the four small ruminant reference assemblies was performed. For each ERV insertion, excluding solo-LTRs, 5kb flanking sequences on each side were extracted. To identify their corresponding positions in the other species, the flanking sequences were aligned to the other assemblies using Minimap2 (version 2.26) (61). If the 5’ and 3’ flanking sequences matched in the same genomic region (within a 50kb interval), and if the interval between them corresponded to an annotated ERV of the same family, the insertion site was considered to be syntenic between the species.

### ERV detection in other ruminant assemblies

The presence of each ERV family was assessed in 20 additional ruminant species (Supplementary Table S1). In addition to the references, several assemblies from *C. hircus (n=4)* and *O. aries (n=23)* from different breeds were analyzed (Supplementary Table S1). The search was performed using BLASTn (47) with default parameters. The hits were filtered using the same criteria as for the reference assembly annotation to respect the 80-80-80 rule (8).

### Statistical analyses

The statistical tests presented in the study were carried out with R (version 4.2.3) (59).

## Results

### Small ruminant species share the same ERV families

Currently, limited information is available regarding endogenous retrovirus (ERV) families in small ruminants, and databases lack consensus sequences for these species. In this study, we describe a total of 23 ERV families present in four reference assemblies of domestic and wild sheep and goats, as well as one assembly for domestic cattle (Fig. 1, Table S1). Among them, 13 were classified as Class I and 10 as Class II endogenous retroviruses according to their relationship with exogenous retroviruses. We used a threshold of at least one conserved protein domain and a minimum of 10 copies to consider a consensus as a valid representative of an ERV family. This could explain why no Class III sequences have been detected; they may be present but as relics. Comparing the different species, 14 families appeared to be shared by the four small ruminants and the cattle species. Six families were only identified in small ruminants (families I-6, I-10, II-5, II-6, II-7, II-9), while three were exclusive to cattle (families II-1, II-4, II-8). However, no family appeared to be specific to either one small ruminant species or genus.

**Fig. 1:**
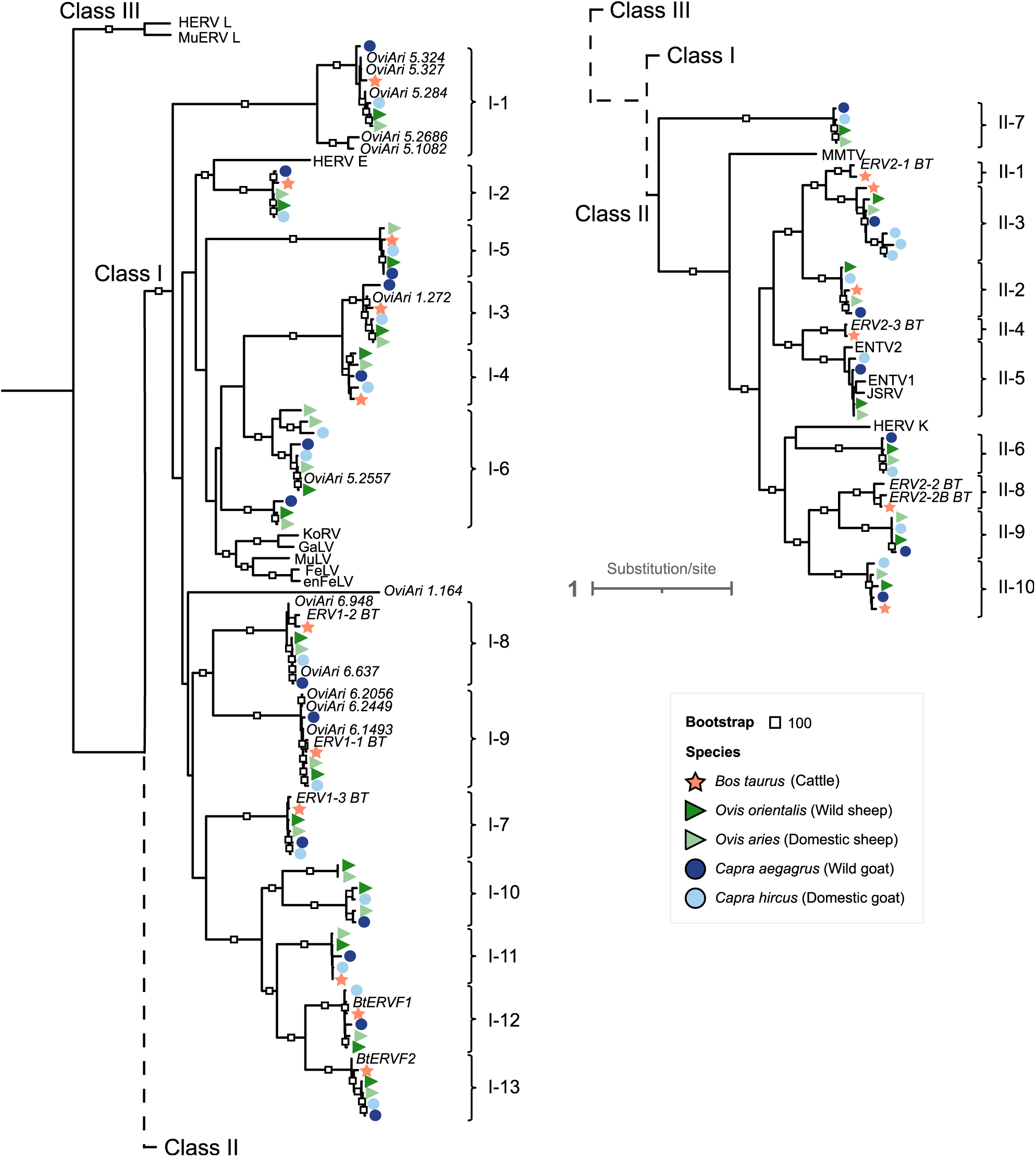
Phylogenetic analysis of the ERV consensus sequences. Maximum Likelihood phylogenetic tree reconstructed from the alignment of the nucleotide internal parts (without LTR) of the consensus sequences generated for the five reference assemblies (represented as colored symbols). Publicly available sequences of endogenous and exogenous retroviruses are indicated by their acronyms, in addition to sheep and cattle Repbase ERV references (in italic) were also included. The ERV family’s names include their classification in Class I or II along with an arbitrary attributed number and are indicated next to each consensus sequence cluster.

More families but also more consensus sequences were identified in small ruminants compared to the cattle, in which only 17 consensus were reconstructed versus 21 to 23 in small ruminants. Although sheep and goats overall shared the same ERV families, the number of consensus sequences between these small ruminant species differed. This variation can be explained by the identification of several consensus sequences for both families I-6 and II-3 that correspond to sub-families (Fig. 1). The genomic structure and the sequences of the consensus also differed between the species and ERV families (Fig. S8, Fig. S9). While seven families (I-6, I-9, I-12, I-13, II-5, II-6, II-7) are represented by consensus sequences with intact coding sequences, the families I-1 and I-5 are represented by less complete consensus with only parts of the retroviral genes. Moreover, some internal consensus sequences were associated to multiple LTR consensus sequences going up to five LTR consensus associated to a single internal consensus for family I-1 in the wild sheep thus reflecting the evolutionary history complexity of some families (Table S6).

The consensus sequences generated in this study were compared to those present in Repbase (Fig. S2, Fig. S3). We successfully recovered all the cattle reference families from Repbase but we also discovered nine new bovine families shared with the small ruminants (families I-1, I-2, I-3, I-4, I-5, I-11, II-2, II-3, II-10). Notably, all the Class II cattle Repbase consensus corresponded to families exclusively found in cattle. For small ruminants, we also identified consensus corresponding to the Repbase references except for OviAri_1.164 and OviAri_5.2686 and we described 15 new small ruminant families. However, we observed redundancy among the Repbase consensus with multiple sequences related to the same family. Sequences identified in this study were characterized with longer retroviral genes and more complete coding sequences than in Repbase which mainly contains uncomplete sheep consensus sequences and wild goat LTR sequences.

### Multiple integration events of ERV families across ruminant evolution

Considering the diversity of the observed ERV families, we inferred their approximate integration times during ruminant evolution. With this objective, we looked for the presence of these ERV families in 20 other ruminant species (Fig. 2, Table S2). Three families (I-1, I-5, I-7) were identified as the oldest ones. They were found in all the analyzed species, suggesting that their first integration occurred more than 40 million years ago (Mya) during the Eocene period. The second oldest families (I-2, I-3, I-4, I-8, I-9, I-11, I-12, I-13, II-10) likely integrated during the Oligocene period, between 25 and 40 Mya. The six ERV families specific to the small ruminants (Fig 1) emerged from multiple integration events since the divergence from the *Bovidae* species. Five families (I-6, I-10, II-6, II-7, II-9) were found in both *Antilopinae* and *Caprinae* species indicating a potential first integration between 14 and 18 Mya. Only family II-5 was exclusively found in *Caprinae* species, suggesting an integration between 6 and 11 Mya. Noteworthy, this family exhibits a close relationship with the exogenous retroviruses ENTV and JSRV (Fig. 1).

**Fig. 2:**
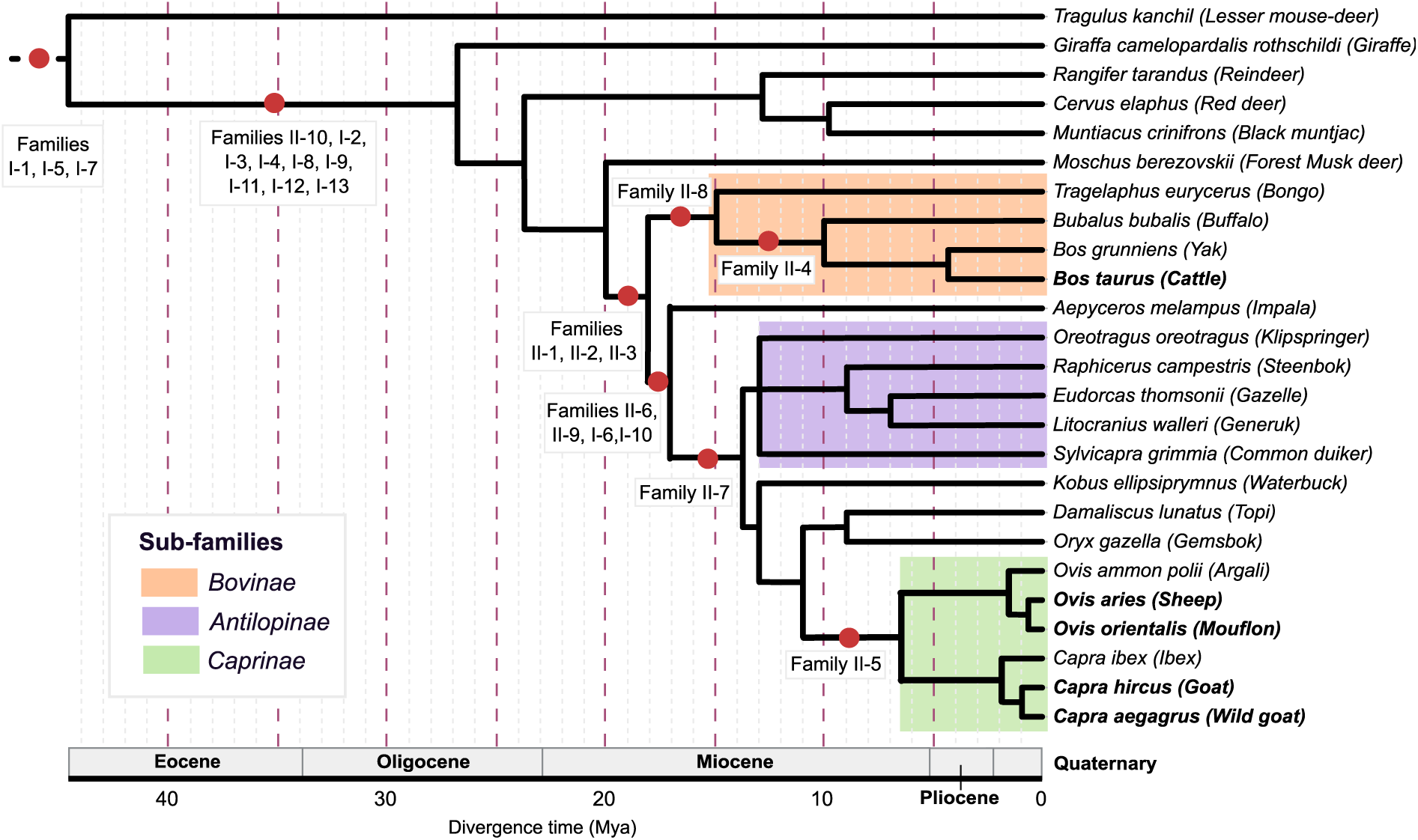
ERV family integration events across ruminant evolution. Each ERV family was detected in both small ruminant and cattle reference assemblies (highlighted in bold) by BLASTn, along with 20 additional ruminant species represented on the tree (see Table S1 for accession numbers). A red dot indicates that the family is present in the following species but absent in the others, suggesting that integration events occurred within the time interval defined by the preceding and subsequent nodes. The phylogeny was produced using TimeTree (97) coupled with divergence time from (98,99).

Among the three families initially exclusively characterized in cattle (Fig. 1), two were present only in *Bovinae* species suggesting integration events from 15 to 18 Mya (family II-8) and 10 to 15 Mya (family II-4), respectively before and after speciation events between bongo and yak on one hand, and buffalo and cattle on the other hand. For the third family, family II-1, no consensus sequence was reconstructed in small ruminant species (Fig 1). However, tracks of copies from this family were found in *Antilopinae* and *Caprinae* species using presence/absence searching analysis suggesting an integration predating *Bovidae* speciation between 17 and 20 Mya (Fig 2). It appears that only very old and degraded copies are present in these species, whereas copies have not disappeared to the same extent in *Bovinae* species, highlighting different ERV family dynamics between species. In summary, ERV families emerged from multiple integration events at different times through the ruminant evolutionary history from over 40 Mya, but no family emerged more recently than 6 Mya, spanning from the end of the Pliocene through the Quaternary periods. Class I families appeared to be older than Class II with only family II-10 older than 25 Mya and families I-6 and I-10 that appeared after *Bovinae* and *Caprinae* speciation.

### Differential insertion dynamics of ERV between species

The annotation of endogenous retroviruses allowed the estimation of their proportion in different genomes, ranging from approximately 18,000 to 25,000 insertions in small ruminants, with a significantly higher number in cattle, where they reach more than 31,000 insertions (Tab 1, Pearson’s χ^2^ = 9460.7, p < 2.2^e^-16). These insertions represent between 0.65% and 1.15% of the different analyzed genomes. Global proportion analysis revealed a predominance of Class I over Class II ERV insertions, except in the bovine assembly where Class II copies outnumbered those from Class I suggesting a difference in insertion activity between cattle and small ruminants. Comparison among small ruminants showed a similar ERV profile between wild and domestic sheep with a significant over-representation of Class I insertions and an under-representation of Class II insertions compared to goats. On the other hand, a higher proportion of ERV copies, especially from Class II, was found in the domestic goat, representing a percentage in the genome twice as high as in the sheep genomes and even higher than in the wild goat genome.

To better understand the differences observed among small ruminant species, the proportion of each ERV family was compared (Fig. 3A). The analysis revealed significant variations in abundance among the different ERV families and across the species for Class I (Pearson’s χ^2^ =3064249, p < 2.2^e^-16) and Class II insertions (Pearson’s χ^2^ = 2971431, p < 2.2^e^-16). The proportion of the different ERV families greatly varied, some being very abundant, with more than 1,000 insertions in the five analyzed assemblies (I-1, I-6, I-7, I-8, I-9, and II-3), while others contain less than 200 insertions (I-11, I-12 and II-9). Among the small ruminant species, distinct quantitative patterns of the different families were observed (Fig. S4). For Class I, the family I-1 showed a significant over-representation in the domestic goat whereas it is under-represented in the wild goat, together with family I-6. For Class II, inter-genus differences between *Ovis* and *Capra* were observed for some families such as II-7 being over-represented in sheep genomes. Species-specific dynamics were observed with families II-3 and II-5 being over-represented in domestic goat (Fig. S4). Furthermore, in cattle, a profile different from the small ruminants was observed, particularly in the context of Class II ERVs (Fig. 3).

**Tab. 1:**
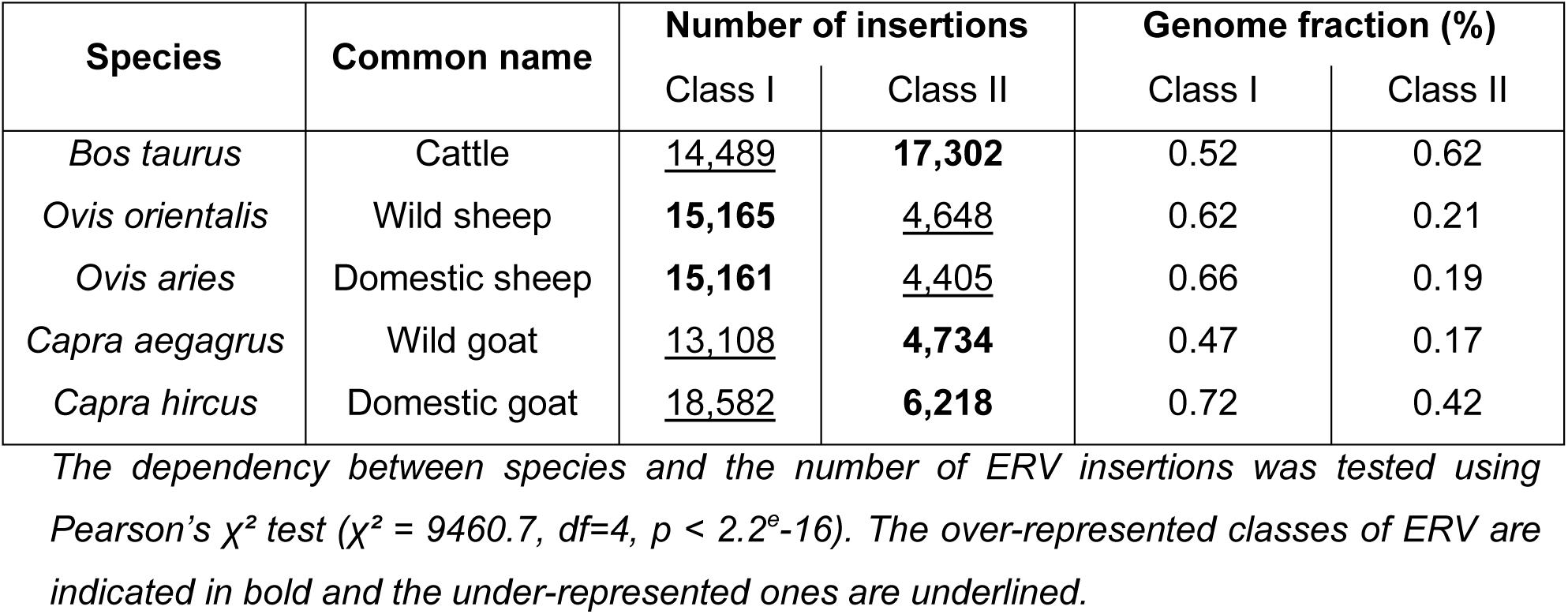
Global ERV proportion in ruminant reference assemblies.

**Fig. 3:**
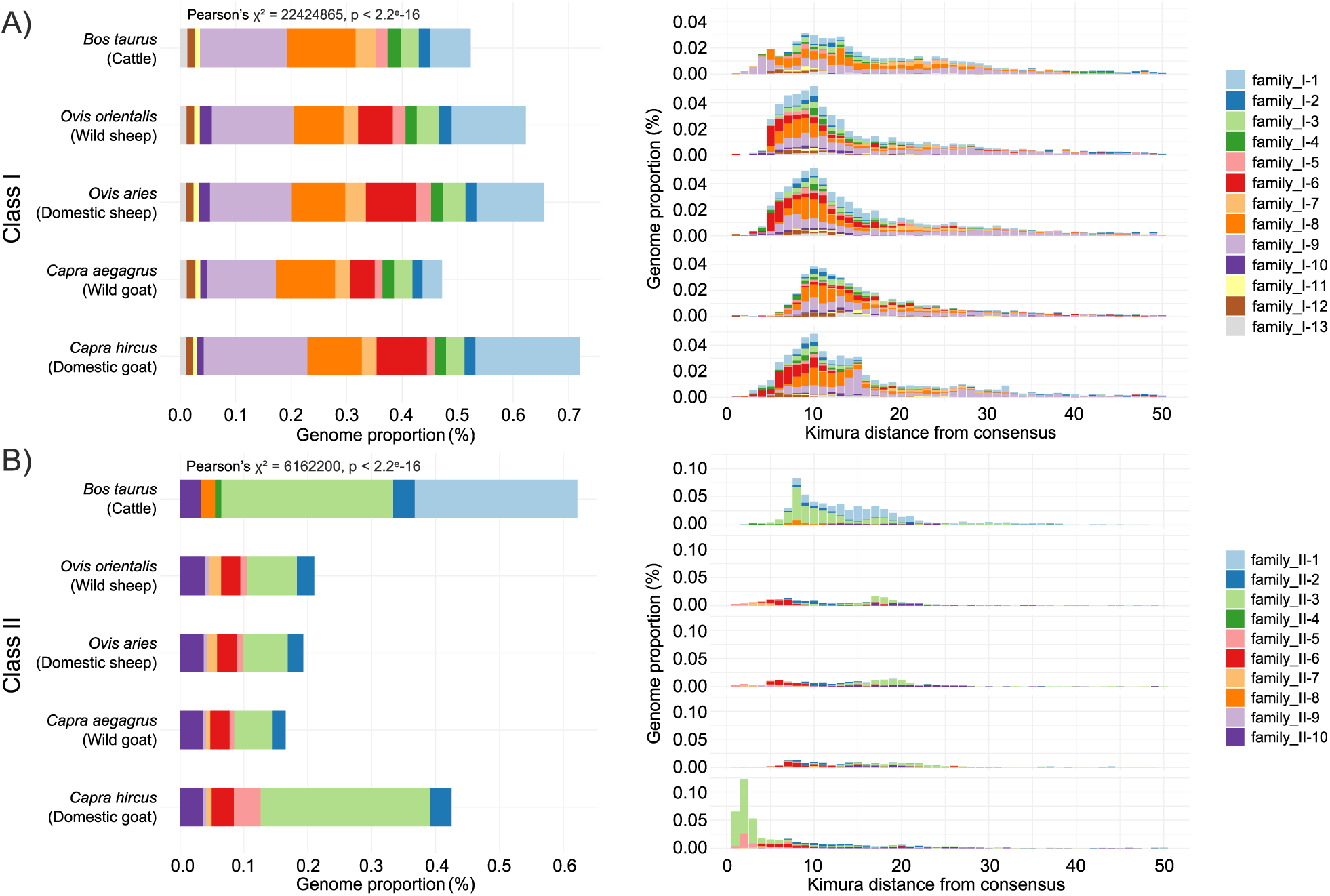
Family proportion and divergence landscapes of ERV in ruminant reference assemblies. Class I and Class II are represented separately in A) and B) panels respectively. The left panels represent ERV family proportions in cattle and small ruminant assemblies. Over- and under-represented families were identified comparing *Caprinae* species (See Pearson’s χ^2^ residuals Fig S4). In the right panels, for each family and Kimura-2 sequence divergence interval, the genome coverage was computed as the percentage of the total ERV insertion length on the total genome length. The divergence distribution was compared between species using the discrete Kolmogorov-Smirnov test and the Benjamini & Hochberg correction (Statistics in Table S4).

Different numbers of insertions were identified between ERV families and species, highlighting the importance of deciphering whether these differences were caused by recent or ancient peaks of transposition activity. Further analyses revealed distinct divergence landscapes between Class I and Class II ERVs (Fig. 3B and Table S4). For Class I, most of the insertions formed a peak with a sequence divergence of around 10% indicating that they are overall no longer active although relatively recent. Only a few copies had a sequence divergence from the consensus close to 0% (Fig. S5) suggesting that they were transposed very recently. Interestingly, these copies were from the families I-6 and I-10 that appeared the more recently in small ruminants and that include copies with intact ORFs for family I-6 suggesting that they could be able to retrotranspose (Table S3).

Class II ERVs exhibited diverse sequence divergence profiles, characterized by multiple peaks suggesting successive waves of ERV activity (Fig. 3B, Fig. S6). Families II-1, II-3, and II-10 displayed peaks of sequence divergence around 20%, consistent with their potentially more ancient emergence among Class II families (Fig. 2). Notably, despite their age, families II-1 and II-3 harbored respectively nine and four copies with intact ORFs in cattle (Table S4), suggesting that these copies could be at the origin of a new activity wave since they are potentially able to retrotranspose. Consistent with their time of appearance after *Bovidae* speciation, families II-5 and II-7 in small ruminants and II-4 and II-8 in cattle showed peaks below 10% of sequence divergence, indicating more recent activity. Family II-5 exhibited recent copies across all small ruminant species but at low frequency. The domestic goat displayed a unique pattern with a significantly high number of recent insertions (divergence <5%) for family II-5 but also for the more ancient family II-3. These results suggest a recent burst of transposition of these families exclusively in the domestic goat. Although most of the divergence distributions were significantly different across species (Table S4) similar distributions were observed for several families including family II-9 among all small ruminants, family II-10 in domestic species and family II-5 among domestic and wild sheep.

### Reactivation of an old family in domestic goat: focus on the family II-3

Copies from the family II-3 were searched in 27 other small ruminant assemblies from different breeds (Fig. 4A, Table S1). The tendency of having more ERV insertions in the domestic goat was not confirmed in other goat assemblies, suggesting that the observed pattern might be specific to the San Clemente breed, used for the ARS1.2 assembly. To investigate why this family contains more copies, we examined the size distribution of the insertions (Fig. 4B). The mean length of the copies was higher in the domestic goat compared to other small ruminant reference assemblies (Wilcoxon test, p-value < 2.2e-16) whereas it was lower in the wild goat (Wilcoxon test, p-value < 2.2e-16). Although the number of solo-LTRs was approximately 2,800 in the four small ruminant species, the number of copies was 10 times higher in the domestic goat, with more than 1,000 copies compared to about a hundred in the others. Moreover, a significant proportion of the domestic goat copies falls within the 2.5 and 6 kb length range, whereas in the other species, only a minority of the copies are truncated. We compared the sequence divergence between the two LTRs of each insertion as a proxy to estimate its insertion date (Fig. S7A). The 5’ and 3’ LTRs were better conserved in the domestic goat insertions with a median of 97.25% sequence identity compared to the wild goat (85.02%, Wilcoxon test, p-adj = 0.00012) and the wild sheep (90.02%, Wilcoxon test, p-adj = 0.02) but not the domestic sheep (96.37%, Wilcoxon test, p-adj = 0.49). This family was however better conserved in cattle as highlighted by the structure of the consensus sequences (Fig. 4E). Indeed, family II-3 consensus sequences was smaller with a length less than 4.5kb in small ruminants compared to 7,973bp in cattle. Consensus sequences from small ruminant lacked parts of their retroviral genes contrary to cattle which had intact ORFs. Among all the copies, five were annotated with full coding capacities in cattle compared to none in small ruminants (Table S3).

**Fig. 4:**
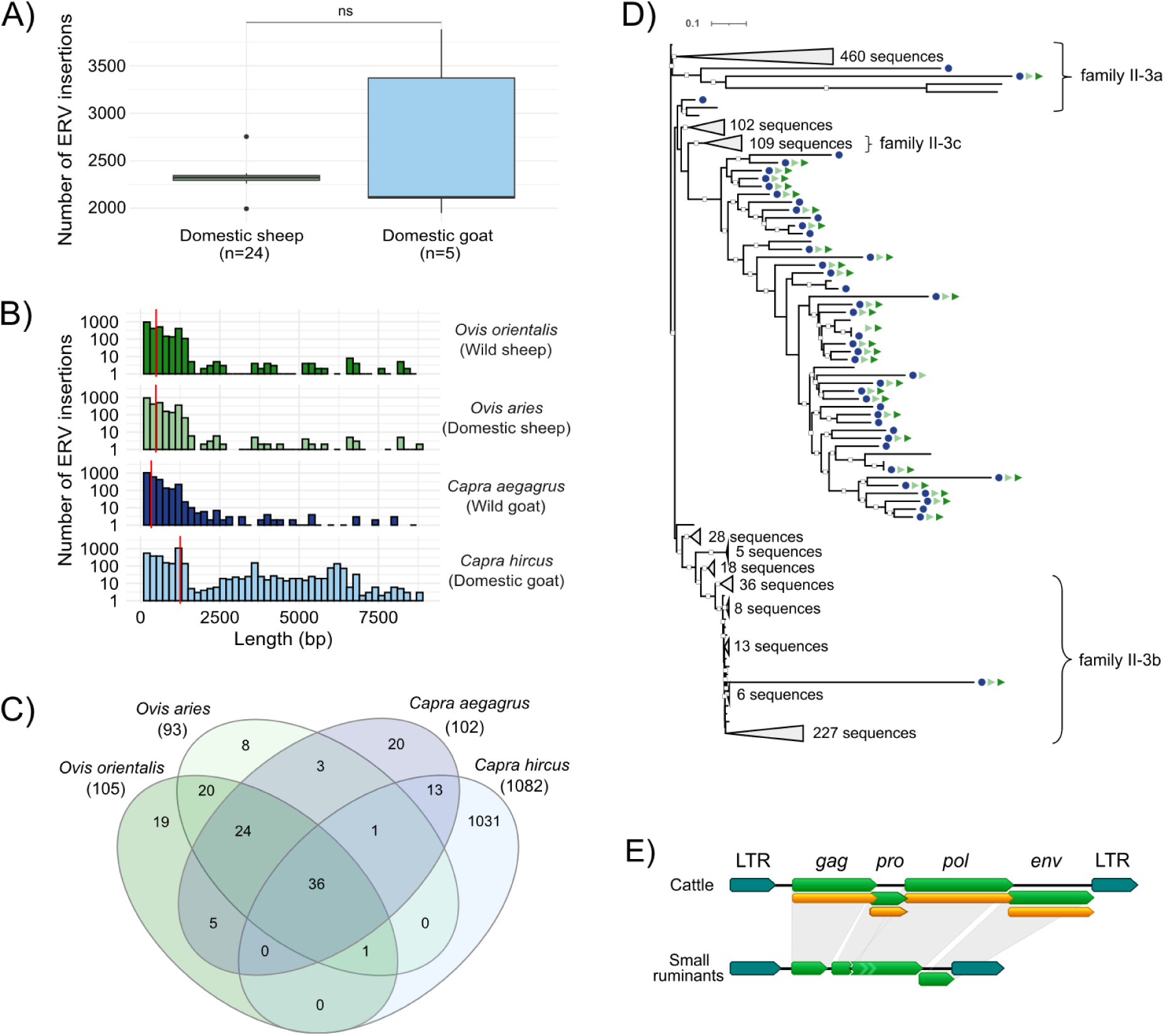
Characteristics of the family II-3 copies in small ruminant genomes. **(A)** Number of family II-3 ERV copies in 29 assemblies from domestic sheep and goat of different breeds (accession numbers in Table S1). **(B)** Length distribution of the ERV insertions for each of the small ruminant reference assembly. The red line indicates the mean length. **(C)** Number of common ERV loci excluding solo-LTRs between the species. **(D)** Phylogenetic tree of the family II-3 copies in *C. hircus* reference genome excluding solo-LTRs. Copies sharing insertion sites with other species are reported with symbols: dark blue circle for wild goat, dark green triangle for wild sheep and soft green circle for domestic sheep. To clarify the tree, nodes including only domestic goat specific insertions without any synteny and nearly identical sequences were collapsed. Three sub-families named as family II-3a, II-3b and II-3c were found in the domestic goat. Copies identified using hits from multiple sub-families thus having ambiguous classification were not included in the tree sub-families. The white squares indicate branches supported by a bootstrap higher than 80. Branch lengths are expressed as the number of substitutions per site. **(E)** Comparison of the family II-3 consensus sequences between domestic cattle and small ruminants. The green boxes represent the retroviral genes and the orange ones the coding sequence (ORF).

We then evaluated if the copies were inserted at the same position in the four small ruminant genomes. Thus, insertion sites of the copies without considering solo-LTRs, were compared between species using a liftover method to convert the ERV positions in other genomes (Fig. 4C). A total of 24 insertion sites were shared between domestic and wild sheep. On the other hand, among the 37 insertions shared between the domestic and wild goat, 36 were also shared with the sheep species suggesting that they integrated before *Ovis* and *Capra* speciation. A low number of insertions were species-specific except for the domestic goat in which 95% of the insertions annotated in its reference genome were not found in the other species. Phylogenetic analysis of the domestic goat copies showed that the goat-specific insertions were closely related with up to 100% sequence identity (Fig. 4D). Almost all of the 36 syntenic and thus older insertions clustered in the same clade. In addition, three sub-families (II-3a, II-3b and II-3c) were identified in the domestic goat and the clades representing sub-families II-3a and II-3b both contained an ancient copy. The presence of a large number of nearly identical sequences, together with the presence of sub-families and closely related ancient insertions, suggests recent reactivations of the family II-3 in goat through different waves of transposition.

### The most recent ERV family is still active in small ruminants: focus on the family II-5

In contrast to the family II-3, domestic goat reference genome appeared to be representative of the *C. hircus* species for the family II-5, as evidenced by a significantly higher number of family II-5 insertions in several goat breeds compared to sheep (Fig. 5A, Wilcoxon test, p-value = 0.00058). The median number of insertions in goats was 223, whereas it was 98 in sheep assemblies, although the insertion numbers varied more in goats than sheep, ranging from 118 to 328 in goats to 66 to 108 in sheep. To explore why this family contains more copies in goats than in sheep, we used the same methodology as for family II-3 and examined the size distribution of the insertions (Fig. 5B). While the mean length of the copies was not significantly higher in domestic goats than in other small ruminants, wild goats had significantly shorter copies than the domestic goat (Wilcoxon test, p-adj < 2.2e-16), and both domestic and wild sheep (Wilcoxon tests, p-adj = 1.2e-14 and p-adj = 1.5e-12 respectively) mainly caused by the absence of full-length copies. In contrast, wild and domestic sheep and the domestic goat have 20, 30 and 78 copies respectively with lengths greater than 7,500bp. The median distance between the two LTRs of each insertion was similar among the domestic goat and the domestic and wild sheep with a sequence divergence lower than 1% (Fig. S7B). However, in the wild goat, the LTRs were significantly more divergent than in the domestic goat (99.30%, Wilcoxon test, p-adj = 0.000088) and both the domestic and wild sheep (99.29% and 99.10% respectively, Wilcoxon test, p-adj = 0.00075 and p-adj = 0.00059). This suggests that this family is well conserved in the small ruminants as confirmed by the structure of the consensus sequences that containing intact ORFs. Surprisingly, a very well conserved consensus was obtained for the wild goat whereas only incomplete insertions were annotated. In comparison, the domestic goat and wild sheep harbored respectively twelve and seven full-length copies respectively while only partially conserved copies were annotated in the domestic sheep assembly.

**Fig. 5:**
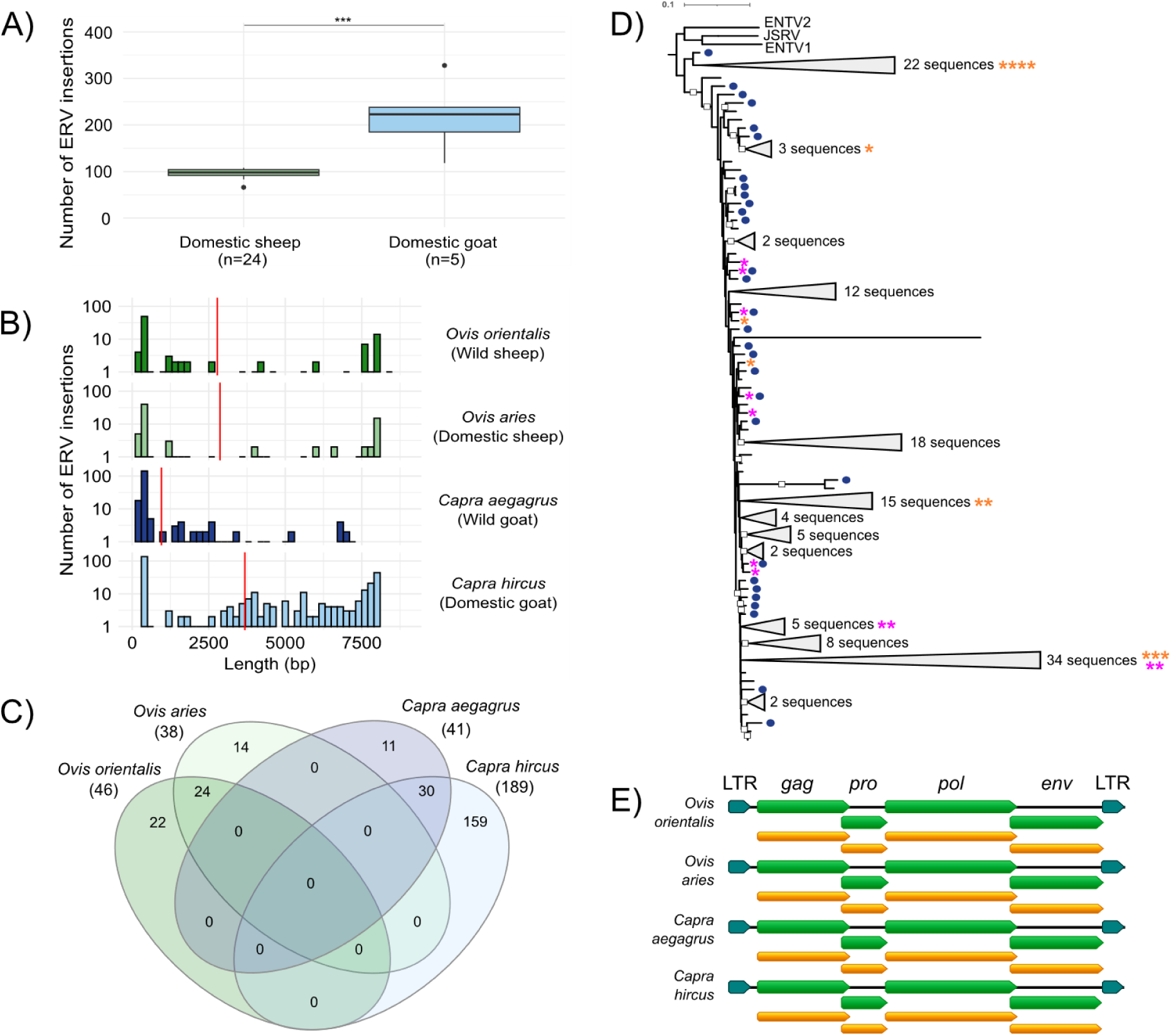
Characteristics of the family II-5 copies in small ruminant genomes. **(A)** Number of family II-5 ERV copies in 29 assemblies from domestic sheep and goat of different breeds (accession numbers in Table S1). **(B)** Length distribution of the ERV insertions for each of the small ruminant reference assembly. The red line indicates the mean length. **(C)** Number of common ERV loci excluding solo-LTRs between the species. **(D)** Phylogenetic tree of the family II-5 copies in *C. hircus* reference genome excluding solo-LTRs. Copies sharing insertion sites with wild goat are represented by dark blue circle. The orange stars represent the insertions with complete *gag*, *pro*, *pol* and *env* ORFs and the pink stars, the copies missing only the complete *env* one. To clarify the tree, nodes including only domestic goat specific insertions without any synteny and nearly identical sequences were collapsed. The white squares indicate the branch supported by a bootstrap of 100. Branch lengths are expressed as the number of substitutions per site. **(E)** Comparison of the family II-5 consensus sequences between small ruminants. The green boxes represent the retroviral genes and the orange ones the coding sequence (ORF).

We described insertions with low sequence divergence compared to their corresponding consensus sequences, with conserved ORFs and with almost identical LTRs indicating that they were very recent copies or that they have been selectively conserved in the small ruminant genomes. To better estimate their integration dates, the different insertion sites of the copies, excluding solo-LTRs, were compared between the species (Fig. 5C). Respectively 24 and 30 insertions were found in the same genomic regions between domestic and wild sheep, or between domestic and wild goats. Regarding the species-specific insertions, 159 insertions were found only in domestic goat representing 84% of the insertions identified in this species. The analysis did not reveal any common insertions between sheep and goats, suggesting that the present copies integrated after the speciation of *Ovis* and *Capra* between 1 and 6 Mya (Fig. 2).

Phylogenetic analysis of the domestic goat copies revealed no discernible clade and indicated close sequence identity between the copies (Fig 5D). Of the identified sequences, 12 contained complete coding sequences, while 11 lacked the *env* ORF. None of the complete coding sequences corresponded to syntenic copies in the wild goats, while three sequences missing only the *env* ORF were in the same genomic region as wild goat copies. Remarkably, the syntenic copies in wild goats showed poor sequence conservation and were all truncated copies highlighting different evolutionary mechanisms involved between wild and domestic goats.

## Discussion

Using a set of bioinformatic tools, we characterized 23 ERV families across five ruminant reference genomes, including cattle and both domestic and wild sheep and goat species. For each ERV family one or more consensus sequences were generated and after multiple filtering steps, were considered as valid representatives of ERV families. We based our manual curation steps on Goubert *et al* (45), but other methods have also been published (62), and several automated pipelines have been developed in recent years (63,64), highlighting the importance of this step and the need to adapt it to specific research questions. Through a comparison with Repbase reference families (53), our results have allowed the refinement of the existing sequences for these species but also the introduction of new reference sequences, particularly for wild and domestic goats, and wild sheep, for which only LTRs or no consensus sequences were previously available.

We identified the presence of both Class I and Class II ERV families in ruminant genomes, in agreement with previous reports on cattle (65–67). While previous studies in sheep have mainly focused on Class II ERV, in particular family II-5 named as enJSRV (35–37,40,41,68–70), nine partial sequences of the *pol* gene from Class I copies have been described (71) corresponding to the families I-4, I-8, I-6 and I-10 in our study. Another study recently described 28 ERV families in *Caprinae* species (72). Three families from their analysis (CapERV-1, CapERV-11, CapERV-26) were not present in our study probably due to the methodology used for *de novo* ERV identification. However, our approach led to the characterization of seven new families including the oldest families (I-1, I-5, I-7) and four additional ones (I-2, I-12, I-13, II-3). Interestingly, two families that we identified with multiple LTR consensus sequences (I-6 and I-10) corresponded to multiple sequences in their analysis confirming the existence of sub-families in these species. Using multiple sequence alignment of the I-6 copies, we were able to distinguish sequence differences between the sub-families in the domestic sheep but it appears that a single internal region coupled with multiple LTR consensus sequences is sufficient to correctly annotate these copies (Fig. S10).

The added value of our study is the comparison of the ERV families between cattle, sheep and goat species. While small ruminants share the same ERV families, differences are observed when compared to cattle. We showed that ruminant ERV families have emerged from multiple integration events across evolution, resulting in ERV families common to both *Bovinae* and *Caprinae*, as well as families specific to each of these *Bovidae* sub-families. Family II-5 was not identified in cattle or other ruminant species but was found exclusively in small ruminants. This contrasts with previous works, where this family was detected in some cattle breeds and in river buffalo (*Bubalus bubalis*) but was absent in others studies (67,73,74). On the other hand, copies from family II-1 seem to be under the influence of the genetic drift and disappearing from most of ruminant species, with the exception of *Bovinae* species where they have been recently described as active (75). These results show that a family can have divergent evolutionary trajectories depending on the breeds but also on the species and underline the diverse dynamics of ERVs influenced by both family and species-specific factors. In addition, dating analyses revealed that ERV Class I families are generally older than Class II families and tend to contain more degraded copies.

By analyzing the number of copies, we estimated that the combined proportion of Class I and II ERVs represents approximately 1% of each ruminant genome, in agreement with previous global studies that estimated their genome fraction at approximately 3% (including Class III copies that represent 2% of the total) (76–78). Our stringent filtering criteria may have led to an underestimation of their genome fraction by missing the most ancient and degraded copies, explaining why we did not encounter any Class III families in our analysis. Even when we focused on the more conserved copies, 84.5% of the identified ERV consisted of solo-LTRs. Overall, few recent insertions were observed, leading to the conclusion that most of the ERV insertions in small ruminants are relics from ancient infection and transposition activity except for families II-3 and II-5 which are still potentially active, especially in the domestic goat.

In the case of family II-3, the domestic goat exhibits a higher number of copies compared to other small ruminants. Common insertion sites were identified across the four species, suggesting integration prior speciation which has since persisted in all genomes. However, many of the copies are goat-specific and located on scaffolds rather than chromosomes. It is important to note that the X chromosome is not assembled at the chromosome level (79) in the goat reference genome, and a proportion of the scaffold-located insertions may in fact originate from this unassembled chromosome. Comparison of the flanking sequences of each goat scaffold-located insertion revealed different insertion sites for some, confirming their existence, but also identical ones for others, which could be the result of satellites from ectopic recombination of retrotransposon LTRs (80–82). Several sub-families have been characterized in the domestic goat genome. One contains almost all syntenic copies together with more recent insertions, and the others two consist mainly of goat-specific elements. This suggests at least two recent bursts of transposition that might have been controlled by a regulatory system (83–87). However, this activity was not detected in the other available goat assemblies and thus would require further investigation using population data.

In contrast, for family II-5, no syntenic insertions were found between *Ovis* and *Capra*, suggesting that all insertions probably occurred after speciation. However, other studies focusing on this ERV family which is closely related to the exogenous JSRV and ENTV, showed at least two insertions (enJSRV-6 and enJSRV-10) shared between sheep and goats suggesting integration before speciation (34,37,43). Identification of the insertion sites of these copies in the available small ruminant genomes using their flanking sequences allowed us to confirm that these two copies are absent from both domestic and wild goat genomes but present in most of sheep genomes (Table S5). These results suggest multiple events of transposition activity and the possible loss of these older insertions in the current goat populations.

Therefore, we characterized two ERV families potentially that may be still active but use different mechanisms for transposition. Family II-5 stands out as the most recent among the families identified, with highly conserved copies across all four small ruminant species. Some of these copies contain intact open reading frames (ORFs), suggesting that they may be capable of autonomous retrotransposition. Furthermore, complete copies could potentially generate virus-like particles capable of infecting other cells and initiating new insertions (34,88–91). However, it is interesting to note that this family is less represented in wild sheep, but contrary to the domestic sheep contain several copies with intact ORFs suggesting possible differences in regulatory systems between species of the same genus. Certain copies from this family have previously been identified as potential invaders of small ruminant genomes (68,92), suggesting that this family has not been completely silenced by transposable element regulatory processes. Other studies have described this family as playing an important role in placental morphogenesis in sheep (93,94) and found expression of these copies in similar tissues in goats (95). However, the lack of syntenic insertions between the two genera raises the question of whether these insertions actually play the same role in the two species, and if the family has been co-opted and is the result of evolutionary convergence. This highlights the need for further investigation into its evolutionary implications including the interplay of the ERVs with their exogenous counterparts JSRV and ENTV (37,43,68). On the other hand, for family II-3, no copy with intact ORFs was found. Our study suggests that duplication events could occur for non-autonomous families, possibly through the recruitment of other proteins via *trans*-complementation. A similar case has been described in cattle for family II-1, known as ERVK[2-1-LTR] (75), which contains copies with intact ORFs in cattle but where most of the *de novo* insertions originate from non-autonomous elements. Retro-elements without LTRs also have been previously described as major players in vertebrate myelination (96).These results further emphasize the complex dynamics of ERV families in ruminants and highlights the diverse strategies employed by these elements in shaping host genomes.

## Conclusions

Our study provides new insights into the evolutionary history of ERV families across ruminant species. Through a comprehensive analysis, we uncovered families with different dynamics including some that remain active within these species. We have generated a robust collection of *de novo* consensus sequences and high-resolution ERV annotations. These results not only deepen our understanding of ERV evolution in ruminants but also provide valuable resources for further exploration into the complex interplay between ERV and host genomes.

## Availability of data and materials

All data generated or analyzed during this study are either publicly available or included in this article and its supplementary information files.

## Acknowledgments

This work was performed using the computing facilities of the CC LBBE/PRABI. We thank F. Arnaud for sharing the unpublished flanking genome sequences of the enJSRV copies reported in (34).

## Founding

This research was funded by the Agence Nationale de la Recherche (ANR), grant ANR-22-CE35-0002-01 and by INRAE GA-SA joint program GoatRetrovirome.

## Contributions

MV performed the analyses and wrote the manuscript. JT, EL and CL conceived the project. JT and EL supervised the work. All participants contributed to the design and implementation of the research, participated to the interpretation of the data and revised the manuscript.

---

**Fig. S1:**
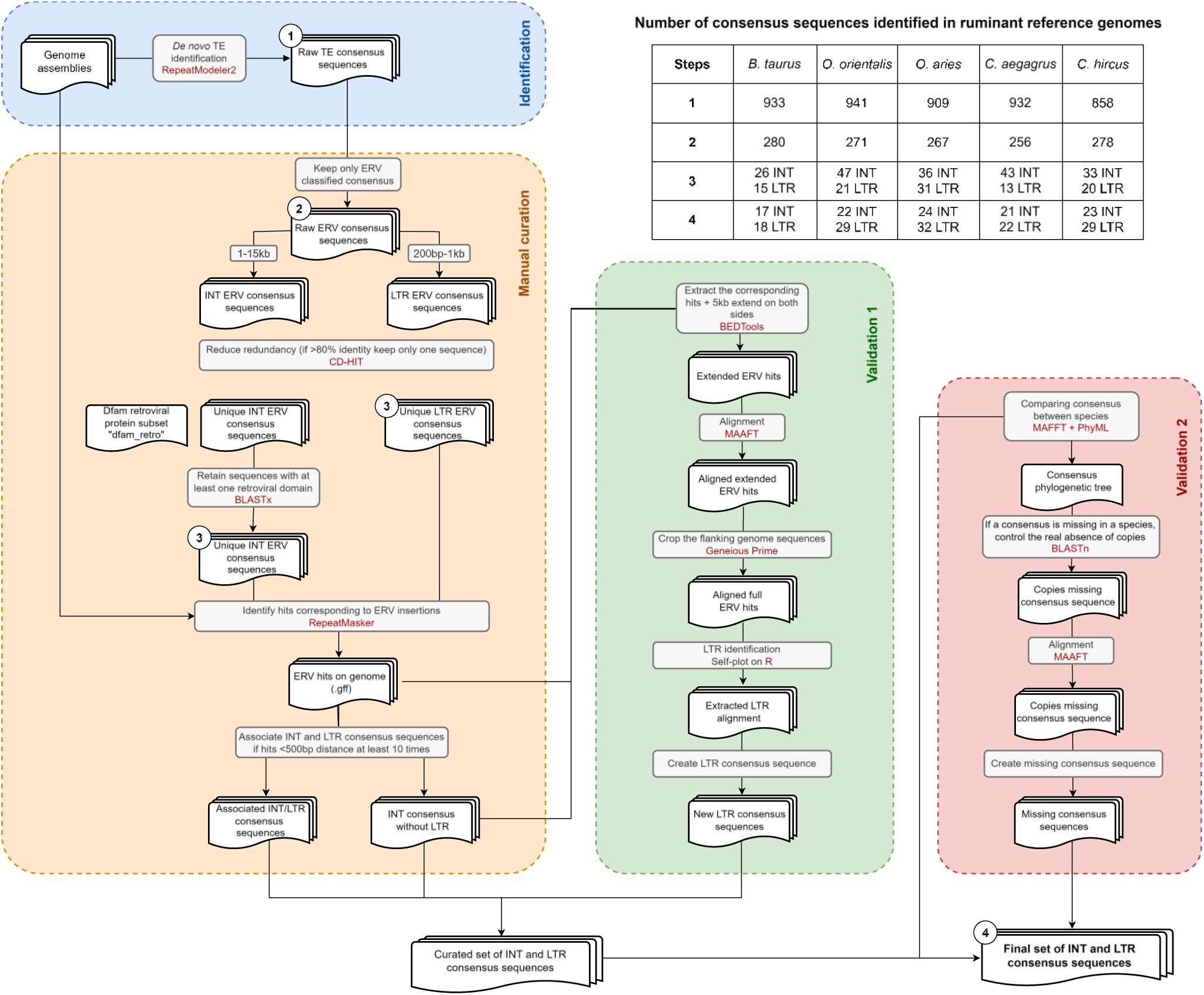
Pipeline used to characterize ERV consensus sequences in ruminant reference genomes. The final set of consensus sequences was obtained after multiple steps going from the *de novo* identification of ERV in the reference assemblies (in blue) to the manual curation and the association between the LTR and internal parts containing the retroviral genes (INT) of the output consensus sequences (in orange) followed by two steps of consensus sequence validation (in green and red). The number of consensus sequences along the pipeline is given in the upper table.

**Fig. S2:**
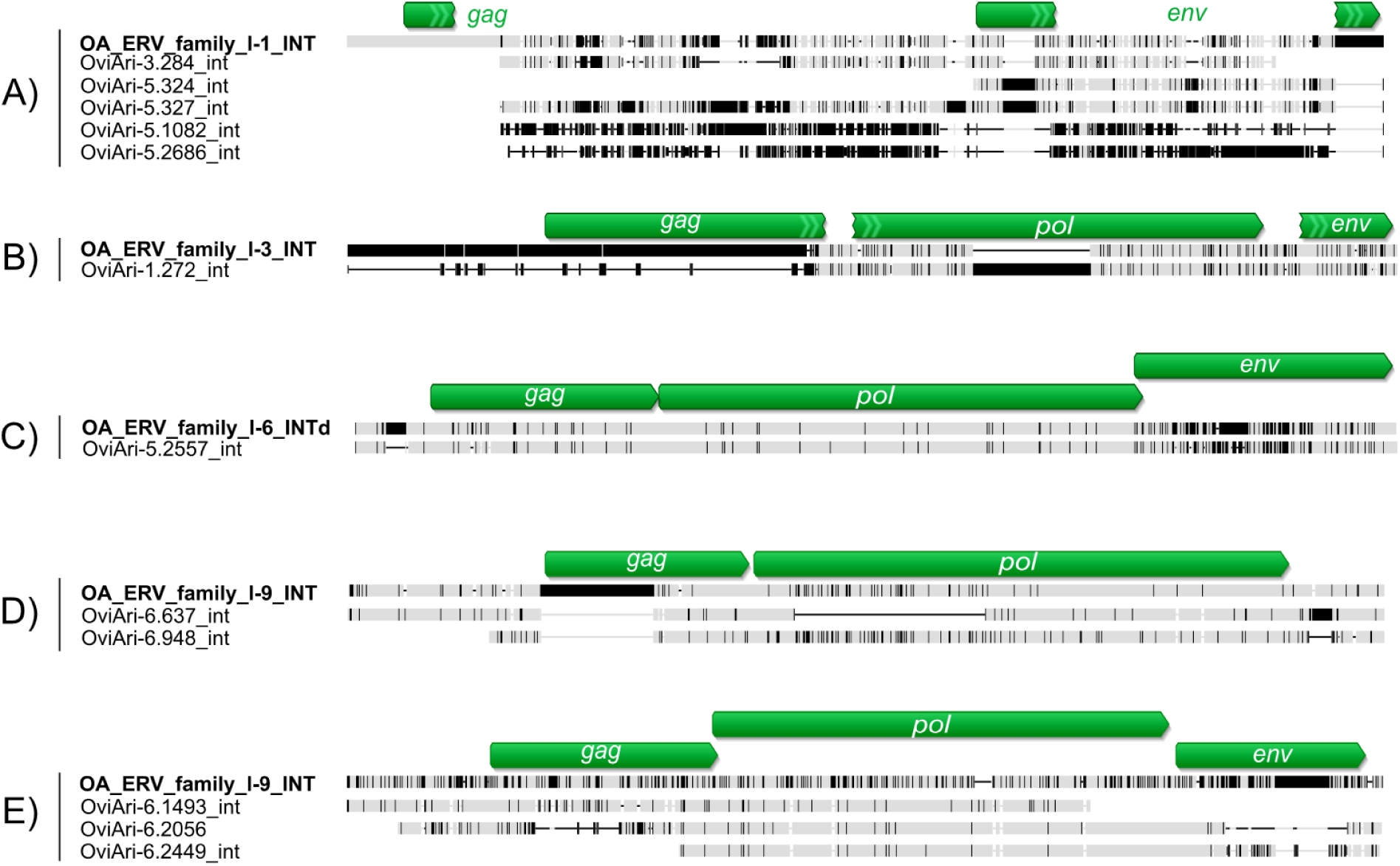
Comparison between Ovis aries ERV Repbase consensus sequences with the ones from this study. The sequences excluding the LTR parts were aligned using MAFFT and visualized with Geneious Prime® (version 2023.0.2). Each panel corresponds to a different ERV family. The consensus sequences characterized in this study are first in raw in bold. The gray boxes indicate similar sequences, the black boxes show sequence differences and the horizontal black lines long deletions. The green arrows represent the retroviral genes.

**Fig. S3:**
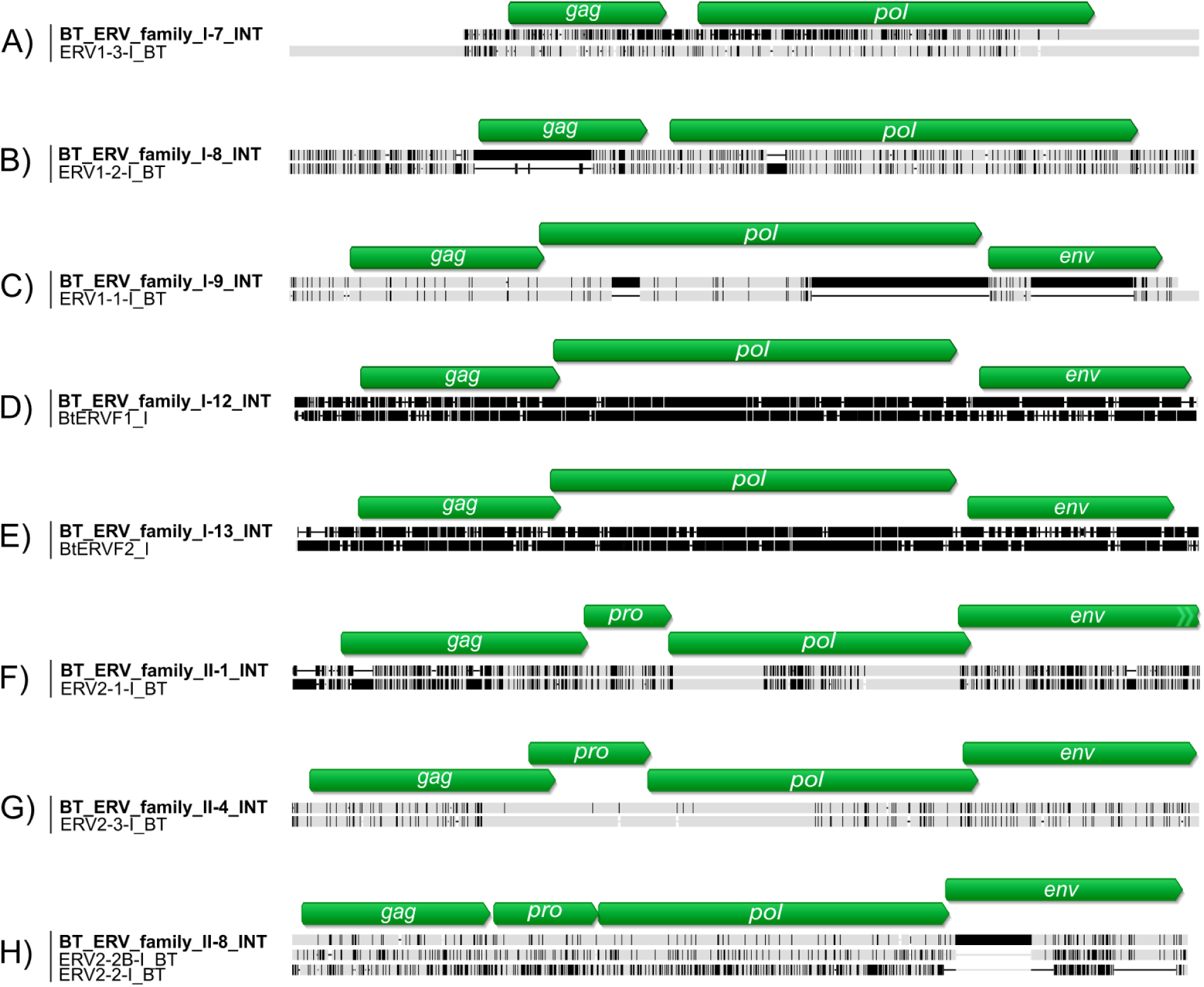
Comparison between Bos taurus ERV Repbase consensus sequences with the ones from this study. The sequences excluding the LTR parts were aligned using MAFFT and visualized with Geneious Prime® (version 2023.0.2). Each panel corresponds to a different ERV family. The consensus sequences characterized in this study are first in raw in bold. The gray boxes indicate similar sequences, the black boxes show sequence differences and the horizontal black lines long deletions. The green arrows represent the retroviral genes.

**Fig. S4:**
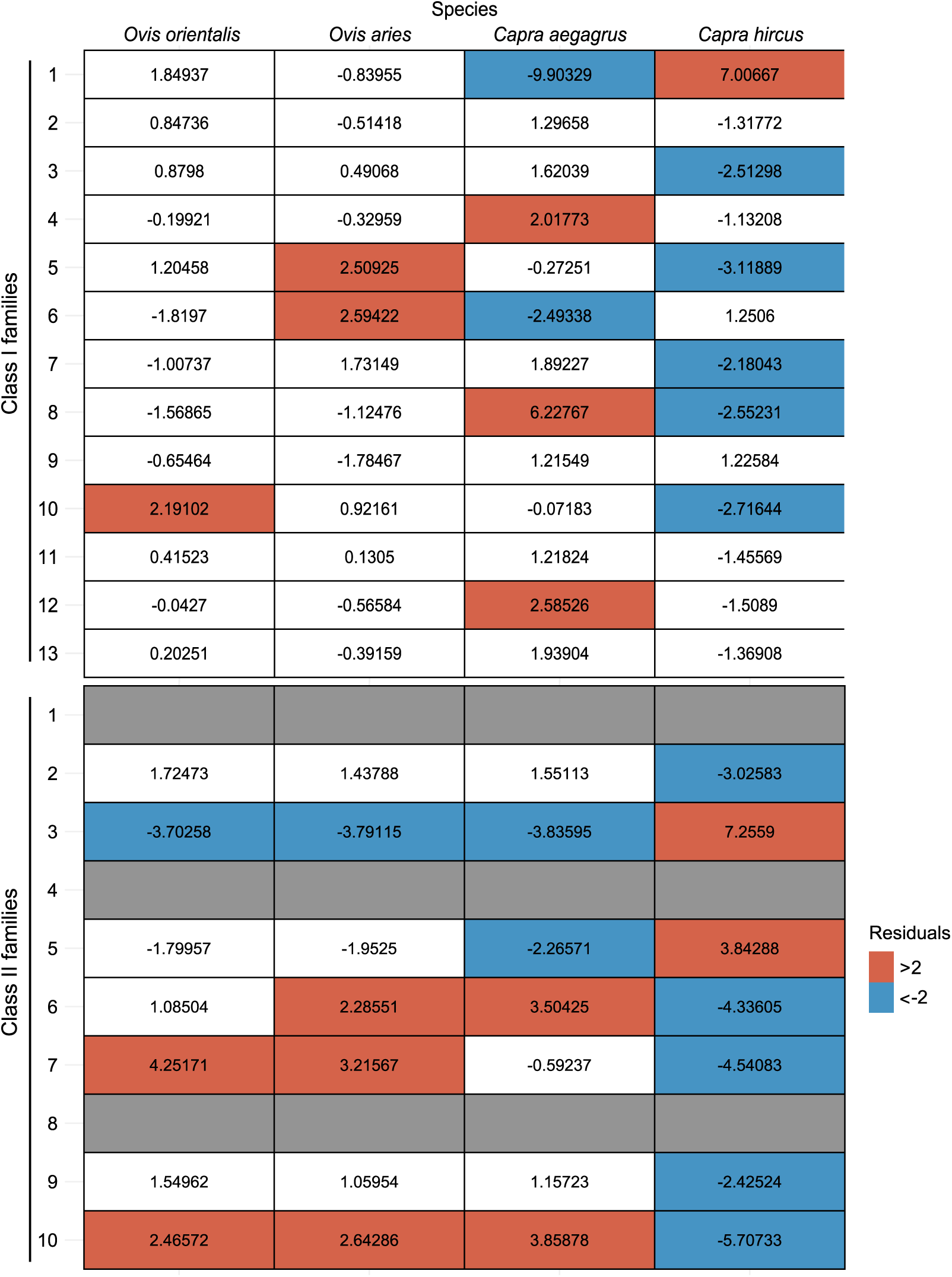
ERV representation between families and small ruminant species. The proportion of each ERV family was compared with a Pearson’s χ^2^ test (Class I: χ^2^ = 3064249, df=36, p < 2.2^e^-16 and Class II: χ^2^ = 2971431, df=18, p < 2.2^e^-16). The χ^2^ residuals are shown here. Families over-represented are in red and the one down-represented in blue. No values were computed for families II-1, II-4 and II-8 only present in cattle but not in small ruminants.

**Fig. S5:**
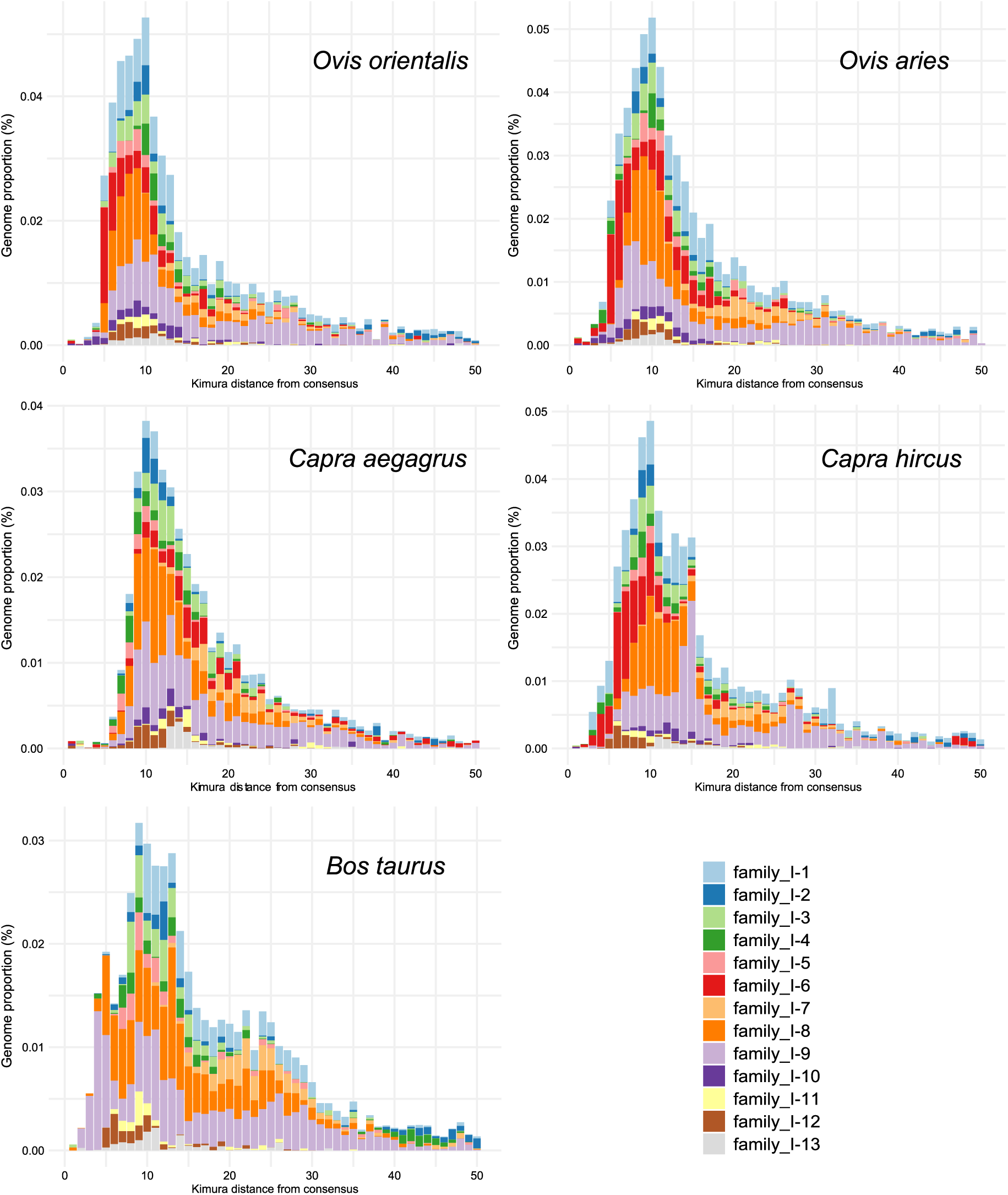
ERV family Class I divergence landscapes in reference ruminant assemblies.

**Fig. S6:**
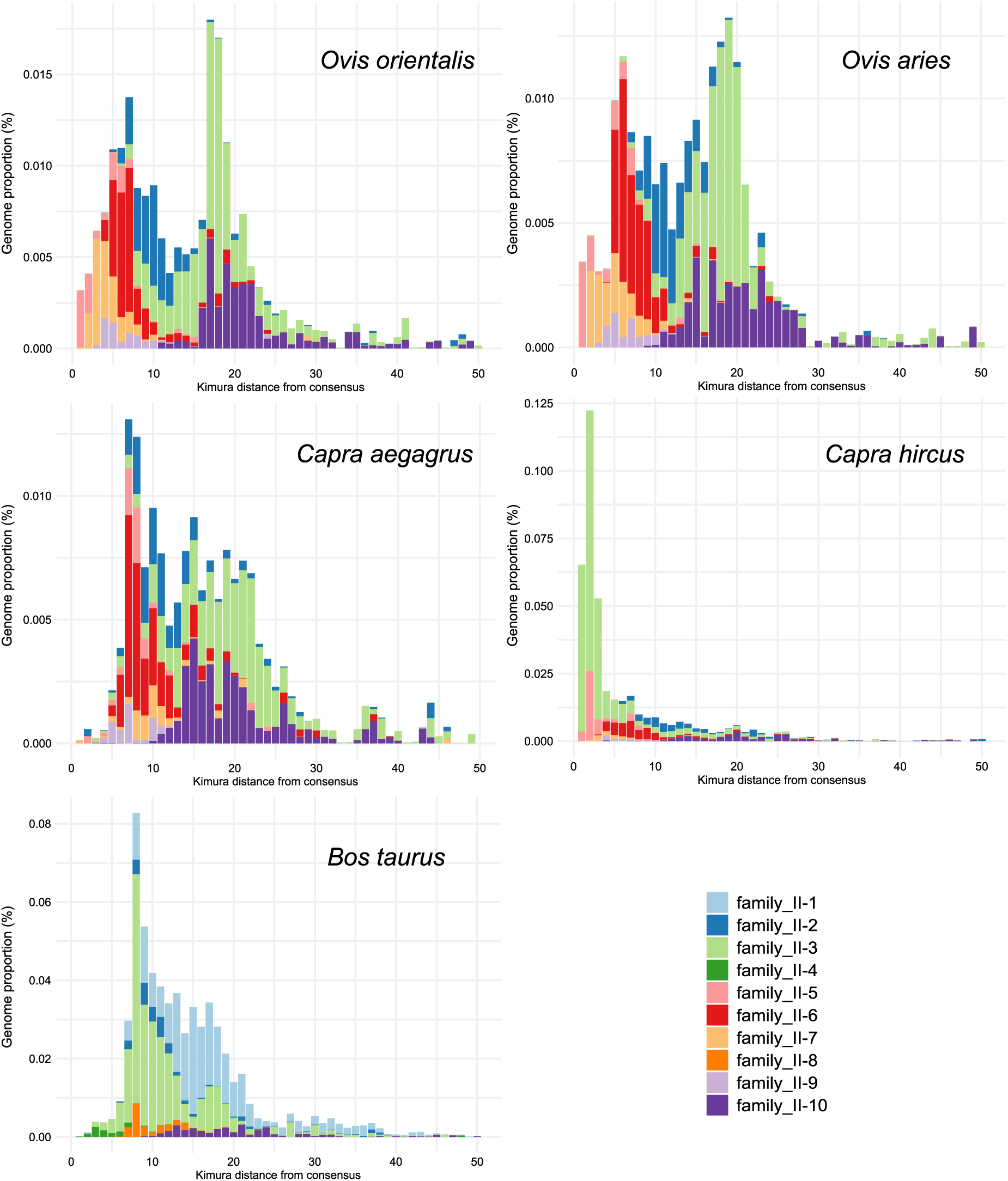
ERV family Class II divergence landscapes in reference ruminant assemblies.

**Fig. S7:**
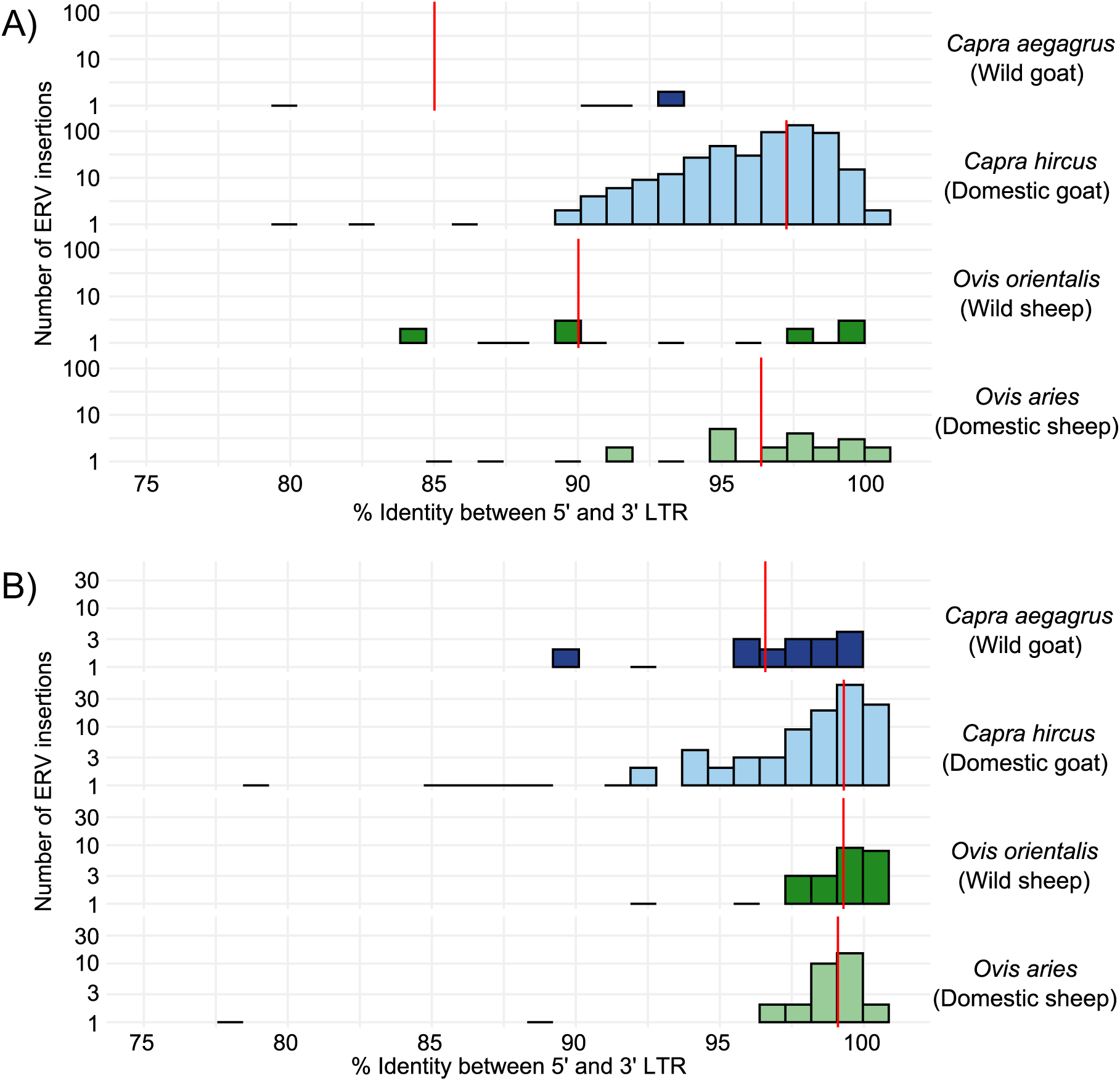
Sequence divergence between ERV family II-3 and II-5 LTRs in small ruminant reference assemblies. The 5’ and 3’ LTRs of each annotated ERV insertion were aligned separately using MAAFT to obtain a pairwise identity score for each LTR pair. Family II-3 is represented in A) and family II-5 in B). For each species, the red line indicates the mean % of identity.

**Fig. S8:**
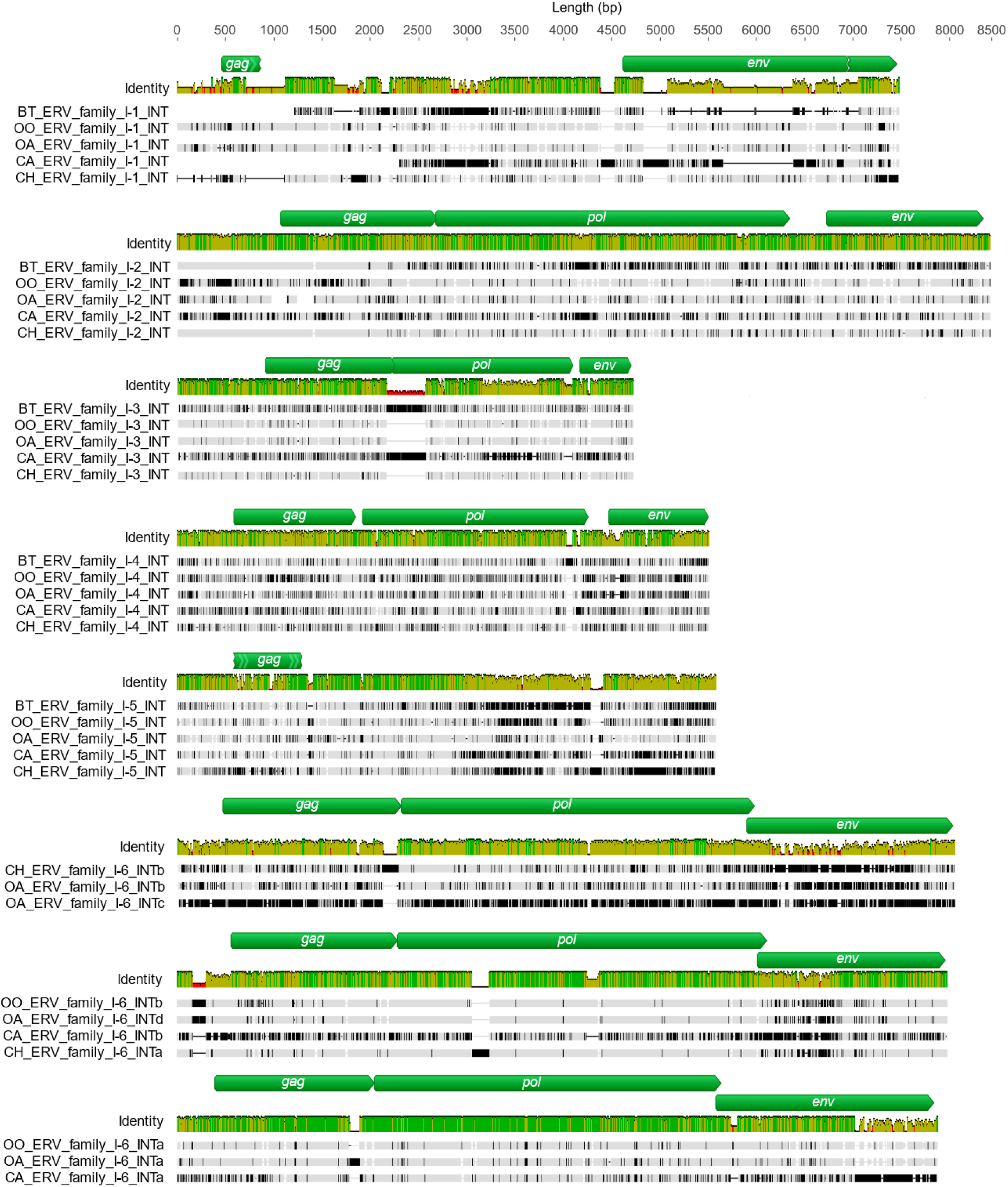
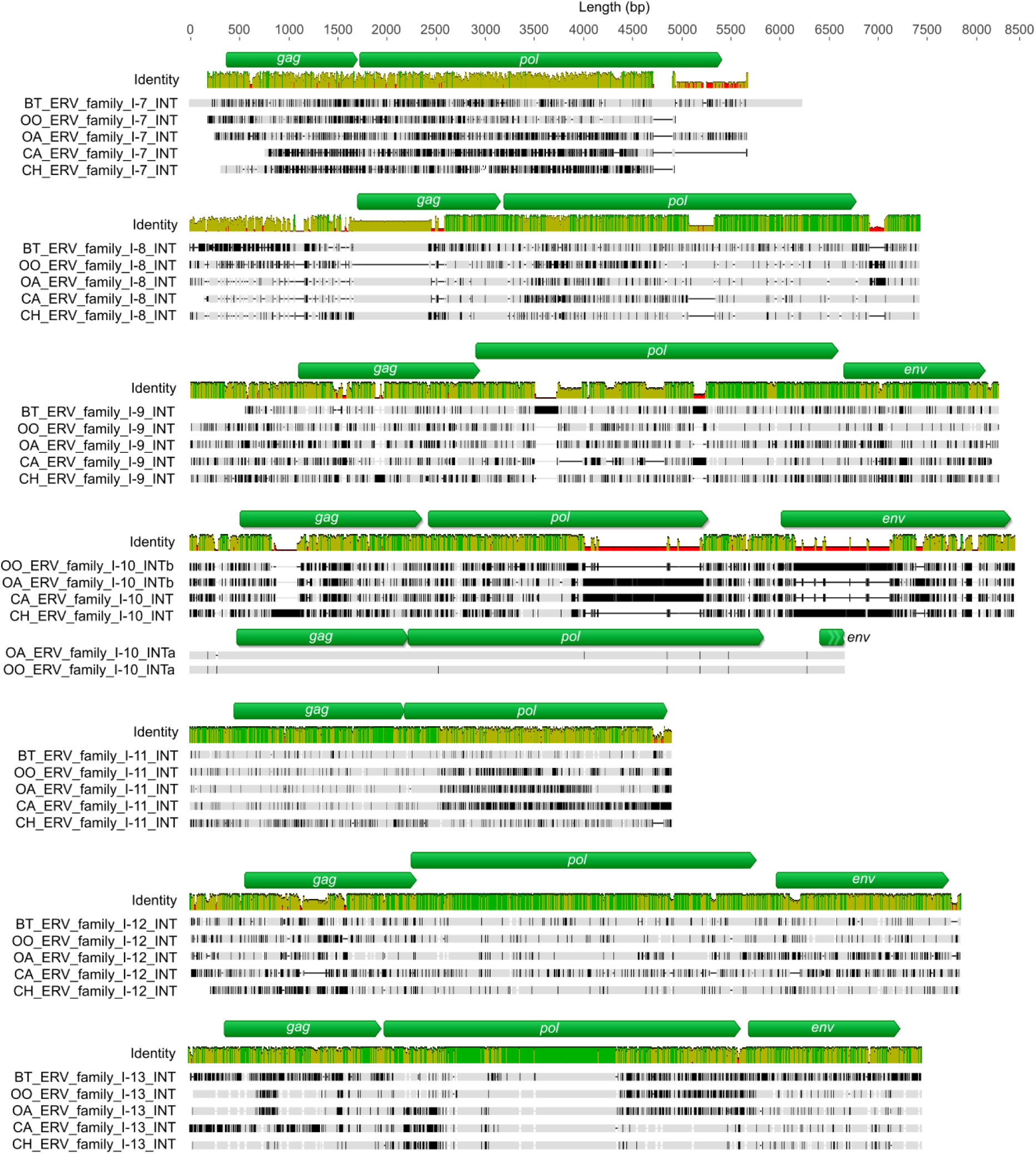
Class I consensus sequences’ genic structure comparison between species. The sequences excluding the LTR parts were aligned using MAFFT and visualized with Geneious Prime® (version 2023.0.2). The gray boxes indicate similar sequences, the black boxes show sequence differences and the horizontal black lines long deletions. The mean pairwise identity over all the sequences for each position is shown above the alignment. Green bars indicate 100% identity, green-brown between 30 and 100% identity and red below 30% identity. The green arrows represent the retroviral genes. Only the internal parts are represented (without the LTR). BT: *B. taurus*; OO: *O. orientalis*; OA: *O. aries*; CA: *C. aegagrus*; CH: *C. hircus*.

**Fig. S9:**
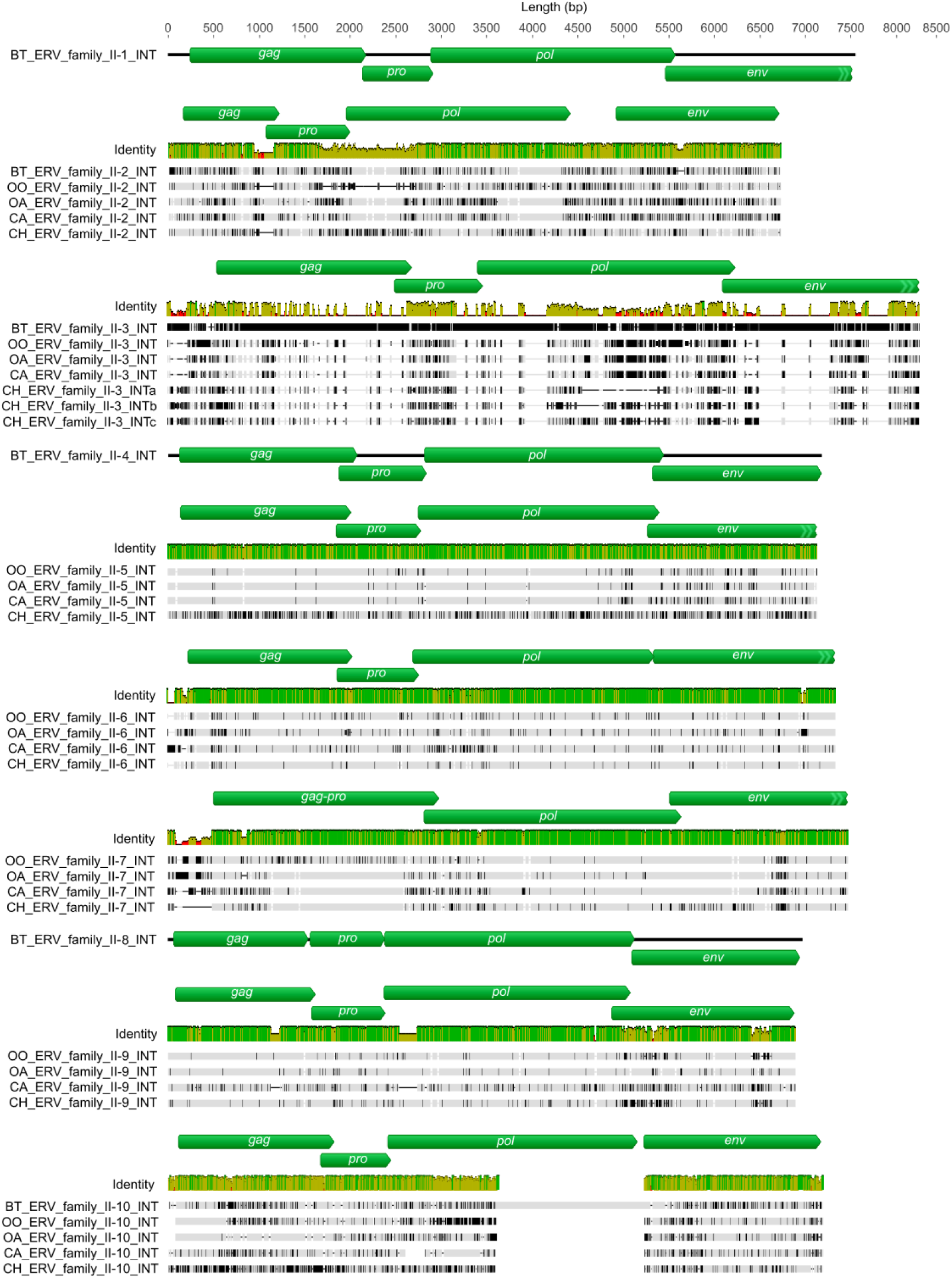
Class II consensus sequences’ genic structure comparison between species. The sequences excluding the LTR parts were aligned using MAFFT and visualized with Geneious Prime® (version 2023.0.2). The gray boxes indicate similar sequences, the black boxes show sequence differences and the horizontal black lines long deletions. The mean pairwise identity over all the sequences for each position is shown above the alignment. Green bars indicate 100% identity, green-brown between 30 and 100% identity and red below 30% identity. The green arrows represent the retroviral genes. Only the internal parts are represented (without the LTR). BT: *B. taurus*; OO: *O. orientalis*; OA: *O. aries*; CA: *C. aegagrus*; CH: *C. hircus*.

**Fig. S10:**
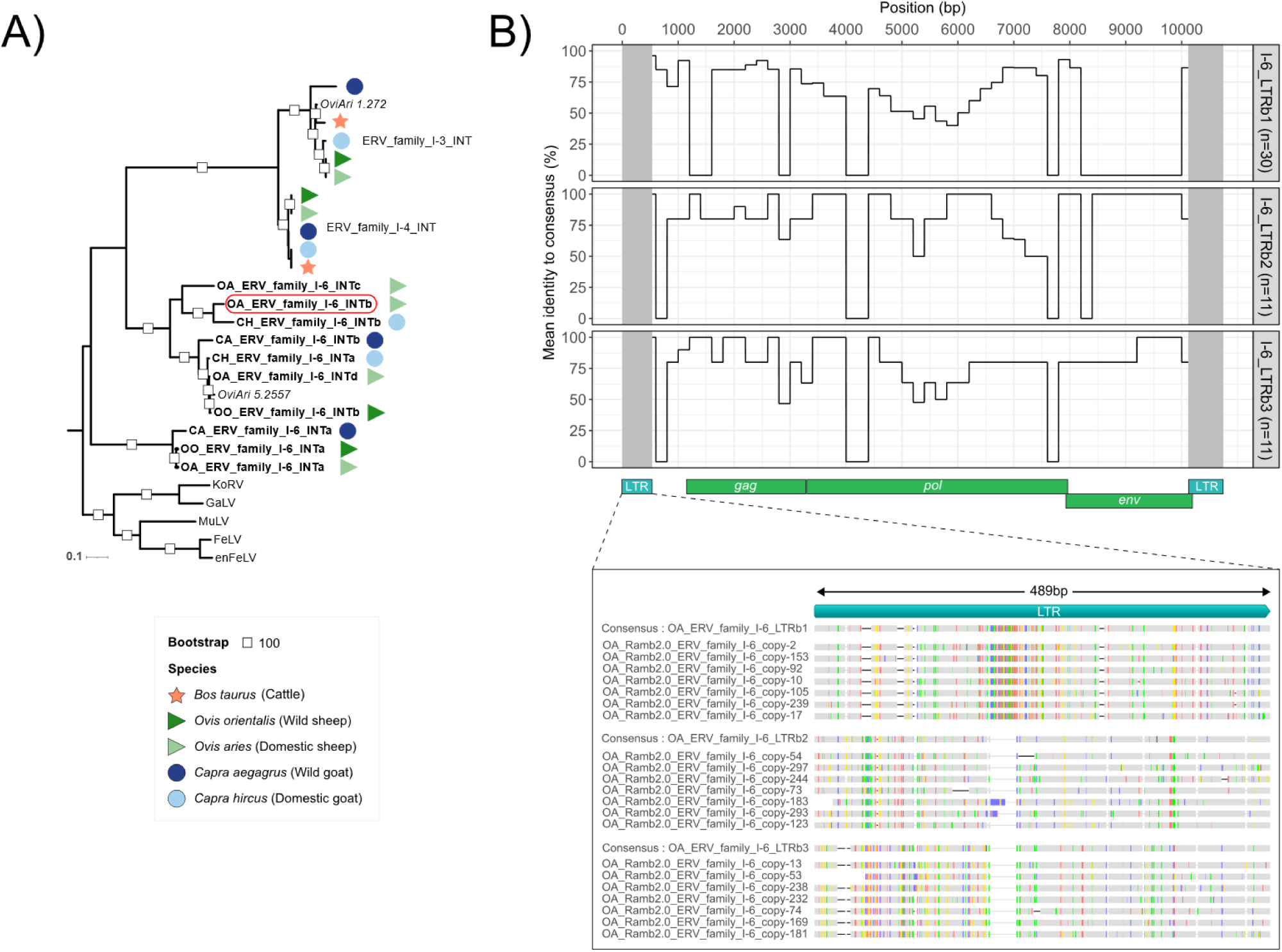
The sub-families of family I-6 in small ruminant genomes. **(A)** ERV family I-6 consensus sequences phylogeny. Maximum Likelihood phylogenetic tree reconstructed from the alignment of the nucleotide internal parts (without LTR) of the consensus sequences generated for the five reference assemblies (represented as colored symbols). Publicly available sequences of endogenous and exogenous retroviruses are indicated by their acronyms, in addition to sheep Repbase ERV references (in italic). Consensus sequences from family I-3 and I-4 were also included. Multiple family I-6 internal sequences were identified in small ruminant (in bold) going up to four sub-families in the domestic sheep (INTa to INTd). Branch lengths are expressed as the number of substitutions per site. **(B)** Multiple LTR sequences of the sub-family I-6b in the domestic sheep genome. A single internal consensus sequence was identified for the sub-family I-6b in the domestic sheep genome, but it was associated to three distinct LTR consensus sequences. All the ERV insertions from sub-family I-6b were aligned to the internal consensus sequence. The 5’ LTR sequences of these ERV insertions were aligned with the different LTR consensus sequences. The first panel displays the average percentage of identity per sub-families at each position of the internal consensus sequence. The second panel presents the alignment of the LTRs, with color-coded boxes representing the sequence differences.

## Additional tables

- Additional_tables_S1-S6.xlsx

**Table S1**: *Ruminant genomes used for ERV detection*.

The five reference assemblies used in this study are indicated in bold. NA: Non-Applicable.

**Table S2:** *Presence/absence of each ERV families in ruminant species*.

**Table S3:** Number of ERV identified in five reference ruminant genomes. NA: Non-Applicable.

**Table S4:** *Kolmogrorov-Smirnov test results on the ERV divergence distribution*.

ERV families with non-significantly different distributions between species (p-adjusted > 0.05) are shown in bold. NA: Non-Applicable.

**Table S5:** *enJSRV insertion sites in small ruminant assemblies*.

**Table S6:** *Number of ERV LTR consensus sequences associated to the INT consensus sequences*.

NA: Non-Applicable.

## Additional files

For all the additional files you can visit the Zenodo archive: https://doi.org/10.5281/zenodo.12188874

- Additional_file_S1.fasta

ERV consensus sequences generated from *Bos taurus* ARS-UCD1.3 assembly.

- Additional_file_S2.fasta

ERV consensus sequences generated from *Ovis orientalis* CAU_Oori_1.0 assembly.

- Additional_file_S3.fasta

ERV consensus sequences generated from *Ovis aries* ARS-UI_Ramb_v2.0 assembly.

- Additional_file_S4.fasta

ERV consensus sequences generated from *Capra aegagrus* CapAeg_1.0 assembly.

- Additional_file_S5.fasta

ERV consensus sequences generated from *Capra hircus* ARS1.2 assembly.

- Additional_file_S6.bed

ERV insertion localization in *Bos taurus* ARS-UCD1.3 assembly.

- Additional_file_S7.bed

ERV insertion localization in *Ovis orientalis* CAU_Oori_1.0 assembly.

- Additional_file_S8.bed

ERV insertion localization in *Ovis aries* ARS-UI_Ramb_v2.0 assembly.

- Additional_file_S9.bed

ERV insertion localization in *Capra aegagrus* CapAeg_1.0 assembly.

- Additional_file_S10.bed

ERV insertion localization in *Capra hircus* ARS1.2 assembly.

- Additional_file_S11.txt

ERV insertion characteristics in *Bos taurus* ARS-UCD1.3 assembly.

- Additional_file_S12.txt

ERV insertion characteristics in *Ovis orientalis* CAU_Oori_1.0 assembly.

- Additional_file_S13.txt

ERV insertion characteristics in *Ovis aries* ARS-UI_Ramb_v2.0 assembly.

- Additional_file_S14.txt

ERV insertion characteristics in *Capra aegagrus* CapAeg_1.0 assembly.

- Additional_file_S15.txt

ERV insertion characteristics in *Capra hircus* ARS1.2 assembly.

## References

1. Weiss RA. The discovery of endogenous retroviruses. Retrovirology. 2006 Oct 3;3(1):67.

2. Stoye JP. Studies of endogenous retroviruses reveal a continuing evolutionary saga. Nat Rev Microbiol. 2012 Jun;10(6):395–406.

3. Feschotte C, Gilbert C. Endogenous viruses: insights into viral evolution and impact on host biology. Nat Rev Genet. 2012 Mar 16;13(4):283–96.

4. Lander ES, Linton LM, Birren B, Nusbaum C, Zody MC, Baldwin J, et al. Initial sequencing and analysis of the human genome. Nature. 2001 Feb 15;409(6822):860–921.

5. Griffiths DJ. Endogenous retroviruses in the human genome sequence. Genome Biol. 2001;2(6):reviews1017.1–reviews1017.5.

6. Mouse Genome Sequencing Consortium, Waterston RH, Lindblad-Toh K, Birney E, Rogers J, Abril JF, et al. Initial sequencing and comparative analysis of the mouse genome. Nature. 2002 Dec 5;420(6915):520–62.

7. Gifford RJ, Blomberg J, Coffin JM, Fan H, Heidmann T, Mayer J, et al. Nomenclature for endogenous retrovirus (ERV) loci. Retrovirology. 2018 Aug 28;15:59.

8. Wicker T, Sabot F, Hua-Van A, Bennetzen JL, Capy P, Chalhoub B, et al. A unified classification system for eukaryotic transposable elements. Nat Rev Genet. 2007 Dec;8(12):973–82.

9. Kapitonov VV, Jurka J. A universal classification of eukaryotic transposable elements implemented in Repbase. Nat Rev Genet. 2008 May;9(5):411–2.

10. Mager DL, Stoye JP. Mammalian Endogenous Retroviruses. Microbiol Spectr. 2015 Feb 5;3(1):10.1128/microbiolspec.mdna3-0009-2014.

11. Zheng J, Wei Y, Han GZ. The diversity and evolution of retroviruses: Perspectives from viral “fossils.” Virol Sin. 2022 Feb 1;37(1):11–8.

12. Sverdlov ED. Perpetually mobile footprints of ancient infections in human genome. FEBS Lett. 1998 May 22;428(1–2):1–6.

13. Miyazawa T, Yoshikawa R, Golder M, Okada M, Stewart H, Palmarini M. Isolation of an Infectious Endogenous Retrovirus in a Proportion of Live Attenuated Vaccines for Pets. J Virol. 2010 Apr;84(7):3690–4.

14. Patience C, Takeuchi Y, Weiss RA. Infection of human cells by an endogenous retrovirus of pigs. Nat Med. 1997 Mar;3(3):282–6.

15. Lavialle C, Cornelis G, Dupressoir A, Esnault C, Heidmann O, Vernochet C, et al. Paleovirology of ‘syncytins’, retroviral env genes exapted for a role in placentation. Philos Trans R Soc B Biol Sci. 2013 Sep 19;368(1626):20120507.

16. Feschotte C. The contribution of transposable elements to the evolution of regulatory networks. Nat Rev Genet. 2008 May;9(5):397–405.

17. Záveský L, Jandáková E, Weinberger V, Minář L, Kohoutová M, Slanař O. Human endogenous retroviruses (HERVs) in breast cancer: altered expression pattern implicates divergent roles in carcinogenesis. Oncology. 2024 Feb 26;

18. Kumar V, McClelland M, Nguyen J, De Robles G, Ittmann M, Castro P, et al. Expression of Endogenous Retroviral RNA in Prostate Tumors has Prognostic Value and Shows Differences among Americans of African Versus European/Middle Eastern Ancestry. Cancers. 2021 Dec 17;13(24):6347.

19. Alldredge J, Kumar V, Nguyen J, Sanders BE, Gomez K, Jayachandran K, et al. Endogenous Retrovirus RNA Expression Differences between Race, Stage and HPV Status Offer Improved Prognostication among Women with Cervical Cancer. Int J Mol Sci. 2023 Jan 12;24(2):1492.

20. Licastro F, Porcellini E. Activation of Endogenous Retrovirus, Brain Infections and Environmental Insults in Neurodegeneration and Alzheimer’s Disease. Int J Mol Sci. 2021 Jul 6;22(14):7263.

21. Römer C. Viruses and Endogenous Retroviruses as Roots for Neuroinflammation and Neurodegenerative Diseases. Front Neurosci. 2021 Mar 12;15:648629.

22. Dhillon P, Mulholland KA, Hu H, Park J, Sheng X, Abedini A, et al. Increased levels of endogenous retroviruses trigger fibroinflammation and play a role in kidney disease development. Nat Commun. 2023 Feb 2;14(1):559.

23. Rangel SC, da Silva MD, da Silva AL, dos Santos J de MB, Neves LM, Pedrosa A, et al. Human endogenous retroviruses and the inflammatory response: A vicious circle associated with health and illness. Front Immunol. 2022 Nov 23;13:1057791.

24. Greenig M. HERVs, immunity, and autoimmunity: understanding the connection. PeerJ. 2019 Apr 5;7:e6711.

25. Vargiu L, Rodriguez-Tomé P, Sperber GO, Cadeddu M, Grandi N, Blikstad V, et al. Classification and characterization of human endogenous retroviruses; mosaic forms are common. Retrovirology. 2016 Jan 22;13:7.

26. Tarlinton RE, Meers J, Young PR. Retroviral invasion of the koala genome. Nature. 2006 Jul;442(7098):79–81.

27. Lillie M, Hill J, Pettersson ME, Jern P. Expansion of a retrovirus lineage in the koala genome. Proc Natl Acad Sci. 2022 Jun 21;119(25):e2201844119.

28. Chiu ES, VandeWoude S. Endogenous Retroviruses Drive Resistance and Promotion of Exogenous Retroviral Homologs. Annu Rev Anim Biosci. 2021 Feb 15;9(Volume 9, 2021):225–48.

29. Mottaghinia S, Stenzel S, Tsangaras K, Nikolaidis N, Laue M, Müller K, et al. A recent gibbon ape leukemia virus germline integration in a rodent from New Guinea. Proc Natl Acad Sci. 2024 Feb 6;121(6):e2220392121.

30. Leroux C, Girard N, Cottin V, Greenland T, Mornex JF, Archer F. Jaagsiekte Sheep Retrovirus (JSRV): from virus to lung cancer in sheep. Vet Res. 2007;38(2):211–28.

31. Leroux C, Mornex JF. Retroviral infections in sheep and the associated diseases. Small Rumin Res. 2008 Apr 1;76(1):68–76.

32. Monot M, Archer F, Gomes M, Mornex JF, Leroux C. Advances in the study of transmissible respiratory tumours in small ruminants. Vet Microbiol. 2015 Dec 14;181(1):170–7.

33. DeMartini JC, Carlson JO, Leroux C, Spencer T, Palmarini M. Endogenous retroviruses related to jaagsiekte sheep retrovirus. Curr Top Microbiol Immunol. 2003;275:117–37.

34. Arnaud F, Caporale M, Varela M, Biek R, Chessa B, Alberti A, et al. A Paradigm for Virus–Host Coevolution: Sequential Counter-Adaptations between Endogenous and Exogenous Retroviruses. PLoS Pathog. 2007 Nov;3(11):e170.

35. Sistiaga-Poveda M, Jugo BM. Evolutionary dynamics of endogenous Jaagsiekte sheep retroviruses proliferation in the domestic sheep, mouflon and Pyrenean chamois. Heredity. 2014 Jun;112(6):571–8.

36. Wang X, Liu S. Endogenous Jaagsiekte sheep retrovirus envelope protein promotes sheep trophoblast cell fusion by activating PKA/MEK/ERK1/2 signaling. Theriogenology. 2022 Nov 1;193:58–67.

37. Armezzani A, Varela M, Spencer TE, Palmarini M, Arnaud F. “Ménage à Trois”: The Evolutionary Interplay between JSRV, enJSRVs and Domestic Sheep. Viruses. 2014 Dec 9;6(12):4926–45.

38. Murcia PR, Arnaud F, Palmarini M. The Transdominant Endogenous Retrovirus enJS56A1 Associates with and Blocks Intracellular Trafficking of Jaagsiekte Sheep Retrovirus Gag. J Virol. 2007 Feb 15;81(4):1762–72.

39. Arnaud F, Murcia PR, Palmarini M. Mechanisms of Late Restriction Induced by an Endogenous Retrovirus. J Virol. 2007 Oct;81(20):11441–51.

40. Armezzani A, Arnaud F, Caporale M, di Meo G, Iannuzzi L, Murgia C, et al. The Signal Peptide of a Recently Integrated Endogenous Sheep Betaretrovirus Envelope Plays a Major Role in Eluding Gag-Mediated Late Restriction. J Virol. 2011 Jul 15;85(14):7118– 28.

41. Viginier B, Dolmazon C, Lantier I, Lantier F, Archer F, Leroux C, et al. Copy Number Variation and Differential Expression of a Protective Endogenous Retrovirus in Sheep. PLOS ONE. 2012 Jul 24;7(7):e41965.

42. Kent M, Moser M, Boman IA, Lindtveit K, Árnyasi M, Sundsaasen KK, et al. Insertion of an endogenous Jaagsiekte sheep retrovirus element into the BCO2 - gene abolishes its function and leads to yellow discoloration of adipose tissue in Norwegian Spælsau (Ovis aries). BMC Genomics. 2021 Jun 30;22(1):492.

43. Cumer T, Pompanon F, Boyer F. Old origin of a protective endogenous retrovirus (enJSRV) in the Ovis genus. Heredity. 2019 Feb;122(2):187–94.

44. Flynn JM, Hubley R, Goubert C, Rosen J, Clark AG, Feschotte C, et al. RepeatModeler2 for automated genomic discovery of transposable element families. Proc Natl Acad Sci. 2020 Apr 28;117(17):9451–7.

45. Goubert C, Craig RJ, Bilat AF, Peona V, Vogan AA, Protasio AV. A beginner’s guide to manual curation of transposable elements. Mob DNA. 2022 Mar 30;13(1):7.

46. Li W, Godzik A. Cd-hit: a fast program for clustering and comparing large sets of protein or nucleotide sequences. Bioinformatics. 2006 Jul 1;22(13):1658–9.

47. Altschul SF, Gish W, Miller W, Myers EW, Lipman DJ. Basic local alignment search tool. J Mol Biol. 1990 Oct 5;215(3):403–10.

48. Storer J, Hubley R, Rosen J, Wheeler TJ, Smit AF. The Dfam community resource of transposable element families, sequence models, and genome annotations. Mob DNA. 2021 Jan 12;12(1):2.

49. Rombel IT, Sykes KF, Rayner S, Johnston SA. ORF-FINDER: a vector for high-throughput gene identification. Gene. 2002 Jan 9;282(1):33–41.

50. Smit, AFA, Hubley, R & Green, P. RepeatMasker Open-4.0. 2013-2015 [Internet]. Available from: https://www.repeatmasker.org

51. Quinlan AR, Hall IM. BEDTools: a flexible suite of utilities for comparing genomic features. Bioinformatics. 2010 Mar 15;26(6):841–2.

52. Katoh K, Rozewicki J, Yamada KD. MAFFT online service: multiple sequence alignment, interactive sequence choice and visualization. Brief Bioinform. 2019 Jul 19;20(4):1160–6.

53. Bao W, Kojima KK, Kohany O. Repbase Update, a database of repetitive elements in eukaryotic genomes. Mob DNA. 2015 Jun 2;6(1):11.

54. Guindon S, Dufayard JF, Lefort V, Anisimova M, Hordijk W, Gascuel O. New Algorithms and Methods to Estimate Maximum-Likelihood Phylogenies: Assessing the Performance of PhyML 3.0. Syst Biol. 2010 May 1;59(3):307–21.

55. Nguyen LT, Schmidt HA, von Haeseler A, Minh BQ. IQ-TREE: a fast and effective stochastic algorithm for estimating maximum-likelihood phylogenies. Mol Biol Evol. 2015 Jan;32(1):268–74.

56. Hoang DT, Chernomor O, von Haeseler A, Minh BQ, Vinh LS. UFBoot2: Improving the Ultrafast Bootstrap Approximation. Mol Biol Evol. 2018 Feb 1;35(2):518–22.

57. Letunic I, Bork P. Interactive Tree Of Life (iTOL) v5: an online tool for phylogenetic tree display and annotation. Nucleic Acids Res. 2021 Jul 2;49(W1):W293–6.

58. Paradis E, Schliep K. ape 5.0: an environment for modern phylogenetics and evolutionary analyses in R. Bioinforma Oxf Engl. 2019 Feb 1;35(3):526–8.

59. R Core Team. R: A Language and Environment for Statistical Computing [Internet]. Vienna, Austria: R Foundation for Statistical Computing; 2023. Available from: https://www.R-project.org/

60. Singh U, Wurtele ES. orfipy: a fast and flexible tool for extracting ORFs. Birol I, editor. Bioinformatics. 2021 Sep 29;37(18):3019–20.

61. Li H. Minimap2: pairwise alignment for nucleotide sequences. Bioinformatics. 2018 Sep 15;34(18):3094–100.

62. Storer J, Hubley R, Rosen J, Smit AFA. Curation guidelines for de novo generated transposable element families. Curr Protoc. 2021 Jun;1(6):e154.

63. Jiangzhao_Qian. qjiangzhao/TEtrimmer [Internet]. 2024. Available from: https://github.com/qjiangzhao/TEtrimmer

64. Orozco-Arias S, Sierra P, Durbin R, González J. MCHelper automatically curates transposable element libraries across species [Internet]. bioRxiv; 2023 [cited 2024 May 21]. p. 2023.10.17.562682. Available from: https://www.biorxiv.org/content/10.1101/2023.10.17.562682v1

65. Xiao R, Park K, Lee H, Kim J, Park C. Identification and Classification of Endogenous Retroviruses in Cattle. J Virol. 2008 Jan;82(1):582–7.

66. Garcia-Etxebarria K, Jugo BM. Evolutionary history of bovine endogenous retroviruses in the Bovidae family. BMC Evol Biol. 2013 Nov 20;13:256.

67. Garcia-Etxebarria K, Jugo BM. Genome-Wide Detection and Characterization of Endogenous Retroviruses in Bos taurus. J Virol. 2010 Oct;84(20):10852–62.

68. Arnaud F, Varela M, Spencer TE, Palmarini M. Coevolution of endogenous Betaretroviruses of sheep and their host. Cell Mol Life Sci CMLS. 2008 Nov;65(21):3422–32.

69. Spencer TE, Palmarini M. Endogenous retroviruses of sheep: a model system for understanding physiological adaptation to an evolving ruminant genome. J Reprod Dev. 2012;58(1):33–7.

70. Qi J wei, Xu M jie, Liu S ying, Zhang Y fei, Liu Y, Zhang Y kun, et al. Identification of Sheep Endogenous Beta-Retroviruses with Uterus-Specific Expression in the Pregnant Mongolian Ewe. J Integr Agric. 2013 May 1;12(5):884–91.

71. Klymiuk N, Müller M, Brem G, Aigner B. Characterization of Endogenous Retroviruses in Sheep. J Virol. 2003 Oct 15;77(20):11268–73.

72. Moawad AS, Wang F, Zheng Y, Chen C, Saleh AA, Hou J, et al. Evolution of Endogenous Retroviruses in the Subfamily of Caprinae. Viruses. 2024 Mar;16(3):398.

73. Morozov VA, Morozov AV, Lagaye S. Endogenous JSRV-like proviruses in domestic cattle: analysis of sequences and transcripts. Virology. 2007 Oct 10;367(1):59–70.

74. Perucatti A, Iannuzzi A, Armezzani A, Palmarini M, Iannuzzi L. Comparative Fluorescence In Situ Hybridization (FISH) Mapping of Twenty-Three Endogenous Jaagsiekte Sheep Retrovirus (enJSRVs) in Sheep (Ovis aries) and River Buffalo (Bubalus bubalis) Chromosomes. Animals. 2022 Jan;12(20):2834.

75. Tang L, Swedlund B, Dupont S, Harland C, Costa Monteiro Moreira G, Durkin K, et al. GWAS reveals determinants of mobilization rate and dynamics of an active endogenous retrovirus of cattle. Nat Commun. 2024 Mar 9;15(1):2154.

76. Ito J, Gifford RJ, Sato K. Retroviruses drive the rapid evolution of mammalian APOBEC3 genes. Proc Natl Acad Sci. 2020 Jan 7;117(1):610–8.

77. Garcia-Etxebarria K, Sistiaga-Poveda M, Jugo BM. Endogenous Retroviruses in Domestic Animals. Curr Genomics. 2014 Aug;15(4):256–65.

78. Hayward A, Cornwallis CK, Jern P. Pan-vertebrate comparative genomics unmasks retrovirus macroevolution. Proc Natl Acad Sci. 2015 Jan 13;112(2):464–9.

79. Bickhart DM, Rosen BD, Koren S, Sayre BL, Hastie AR, Chan S, et al. Single-molecule sequencing and chromatin conformation capture enable de novo reference assembly of the domestic goat genome. Nat Genet. 2017 Apr;49(4):643–50.

80. Dias GB, Svartman M, Delprat A, Ruiz A, Kuhn GCS. Tetris Is a Foldback Transposon that Provided the Building Blocks for an Emerging Satellite DNA of Drosophila virilis. Genome Biol Evol. 2014 May 24;6(6):1302–13.

81. Sharma A, Wolfgruber TK, Presting GG. Tandem repeats derived from centromeric retrotransposons. BMC Genomics. 2013 Mar 4;14:142.

82. Macas J, Koblížková A, Navrátilová A, Neumann P. Hypervariable 3′ UTR region of plant LTR-retrotransposons as a source of novel satellite repeats. Gene. 2009 Dec 15;448(2):198–206.

83. Fultz D, Choudury SG, Slotkin RK. Silencing of active transposable elements in plants. Curr Opin Plant Biol. 2015 Oct 1;27:67–76.

84. Almeida MV, Vernaz G, Putman ALK, Miska EA. Taming transposable elements in vertebrates: from epigenetic silencing to domestication. Trends Genet TIG. 2022 Jun;38(6):529–53.

85. Czech B, Hannon GJ. One Loop to Rule Them All: The Ping-Pong Cycle and piRNA-Guided Silencing. Trends Biochem Sci. 2016 Apr;41(4):324–37.

86. Wang J, Yuan L, Tang J, Liu J, Sun C, Itgen MW, et al. Transposable element and host silencing activity in gigantic genomes. Front Cell Dev Biol. 2023 Feb 24;11:1124374.

87. Deniz Ö, Frost JM, Branco MR. Regulation of transposable elements by DNA modifications. Nat Rev Genet. 2019 Jul;20(7):417–31.

88. Arnaud F, Black SG, Murphy L, Griffiths DJ, Neil SJ, Spencer TE, et al. Interplay between Ovine Bone Marrow Stromal Cell Antigen 2/Tetherin and Endogenous Retroviruses. J Virol. 2010 May;84(9):4415–25.

89. Black SG, Arnaud F, Burghardt RC, Satterfield MC, Fleming JAGW, Long CR, et al. Viral Particles of Endogenous Betaretroviruses Are Released in the Sheep Uterus and Infect the Conceptus Trophectoderm in a Transspecies Embryo Transfer Model. J Virol. 2010 Sep;84(18):9078–85.

90. Fábryová H, Hron T, Kabíčková H, Poss M, Elleder D. Induction and characterization of a replication competent cervid endogenous gammaretrovirus (CrERV) from mule deer cells. Virology. 2015 Nov 1;485:96–103.

91. Preuss T, Fischer N, Boller K, Tönjes RR. Isolation and Characterization of an Infectious Replication-Competent Molecular Clone of Ecotropic Porcine Endogenous Retrovirus Class C. J Virol. 2006 Oct 15;80(20):10258–61.

92. Wang J, Han GZ. Genome mining shows that retroviruses are pervasively invading vertebrate genomes. Nat Commun. 2023 Aug 17;14(1):4968.

93. Dunlap KA, Palmarini M, Varela M, Burghardt RC, Hayashi K, Farmer JL, et al. Endogenous retroviruses regulate periimplantation placental growth and differentiation. Proc Natl Acad Sci U S A. 2006 Sep 26;103(39):14390–5.

94. Cornelis G, Heidmann O, Degrelle SA, Vernochet C, Lavialle C, Letzelter C, et al. Captured retroviral envelope syncytin gene associated with the unique placental structure of higher ruminants. Proc Natl Acad Sci. 2013 Feb 26;110(9):E828–37.

95. Caporale M, Martineau H, De las Heras M, Murgia C, Huang R, Centorame P, et al. Host Species Barriers to Jaagsiekte Sheep Retrovirus Replication and Carcinogenesis. J Virol. 2013 Oct;87(19):10752–62.

96. Ghosh T, Almeida RG, Zhao C, Mannioui A, Martin E, Fleet A, et al. A retroviral link to vertebrate myelination through retrotransposon-RNA-mediated control of myelin gene expression. Cell. 2024 Feb 15;187(4):814–830.e23.

97. Kumar S, Suleski M, Craig JM, Kasprowicz AE, Sanderford M, Li M, et al. TimeTree 5: An Expanded Resource for Species Divergence Times. Mol Biol Evol. 2022 Aug 1;39(8):msac174.

98. Chen L, Qiu Q, Jiang Y, Wang K, Lin Z, Li Z, et al. Large-scale ruminant genome sequencing provides insights into their evolution and distinct traits. Science. 2019 Jun 21;364(6446):eaav6202.

99. Chen ZH, Xu YX, Xie XL, Wang DF, Aguilar-Gómez D, Liu GJ, et al. Whole-genome sequence analysis unveils different origins of European and Asiatic mouflon and domestication-related genes in sheep. Commun Biol. 2021 Nov 18;4(1):1–15.

